# Integrating Long-Read Structural Variant Analysis with single-nucleus RNA-seq to Elucidate Gene Expression Effects in Disease

**DOI:** 10.64898/2026.03.20.713192

**Authors:** Kwanho Kim, Zechuan Lin, Sean K. Simmons, Jacob Parker, Matthew Kearney, Zhixiang Liao, Nathan Haywood, Jixiang Zhang, Madison P. Cline, Idil Tuncali, Monika Sharma, Geidy E. Serrano, Thomas G. Beach, Xianjun Dong, Victoria Popic, Clemens R. Scherzer, Joshua Z. Levin

**Affiliations:** Aligning Science Across Parkinson’s (ASAP) Collaborative Research Network, Chevy Chase, MD 20815, USA; Stanley Center for Psychiatric Research, Broad Institute of MIT and Harvard, Cambridge, MA 02142, USA; Stephen & Denise Adams Center for Parkinson’s Disease Research of Yale School of Medicine, New Haven, CT 06510, USA; APDA Center for Parkinson Precision Medicine, Yale School of Medicine, New Haven, CT 06510, USA; Department of Neurology, Yale School of Medicine, New Haven, CT 06510, USA; Department of Genetics, Yale, New Haven, CT 06510, USA; Department of Neuroscience, Yale, New Haven, CT 06510, USA; Precision Neurology Program of Brigham & Women’s Hospital, Harvard Medical School, Boston, MA 02115, USA; Banner Sun Health Research Institute, Sun City, AZ 85351, USA; Broad Clinical Labs, Broad Institute of MIT and Harvard, Cambridge, MA 02142, USA

## Abstract

Structural variants (SVs) are a major source of genetic diversity, yet how they impact cell types in complex brain diseases remains largely unexplored, partially due to limitations of short-read sequencing. Here, we addressed this fundamental question in Parkinson’s disease (PD). generating long-read whole-genome sequencing (WGS) data for 100 post-mortem brain samples from a PD cohort, constructing a high-confidence catalog of 74,552 SVs. To resolve their functional impact, we integrated single-nucleus RNA-sequencing data from two brain regions from the same samples and focused functional analyses on SVs proximal to genes previously nominated as *cis*-regulated, potential causal targets of PD-associated GWAS loci. Using expression quantitative trait locus and allele-specific expression analyses, we uncovered SVs significantly associated with expression in specific cell types as well as effects shared across cell types. This study demonstrates the power of uniting long-read WGS with transcriptomics to uncover SVs underlying complex disease architecture with cell type resolution.

## Introduction

Structural variants (SVs)—a class of variation that includes insertions, deletions, duplications, and more complex rearrangements—are a major source of genetic diversity and have a profound impact on gene function and regulation^1–3^. While resolving the full spectrum of SVs is still a challenge, the advent of long-read sequencing of human genomes^4,5^ and subsequently high-fidelity (HiFi) long reads with >99.8% accuracy was a technological turning point^6^. With longer reads that span repetitive genomic regions more accurately, we can now generate a more comprehensive map of genetic variation^1,7–10^, a crucial capability given that SVs are disproportionately enriched in and mediated by repetitive DNA^10^. More accurate information about SVs from long-read sequencing enables better downstream functional analyses that assess association of SVs to gene expression (expression quantitative trait locus (eQTL)) and to allele-specific expression (ASE, allelic imbalance)^11,12^. Leveraging the current state-of-the-art technology, this approach enables the creation of a high-confidence catalog of SVs, which is essential for investigating their contribution to complex diseases.

Parkinson’s disease (PD) is a progressive neurodegenerative disorder with a substantial genetic component^13,14^. Landmark genome-wide association studies (GWAS) have been instrumental in mapping its genetic landscape, successfully identifying 78 independent risk loci (90 GWAS peaks) associated with the disease^15^. However, these studies, which primarily survey single nucleotide polymorphisms (SNPs), have not included SVs, which are generally more likely than SNPs to be mapped as causal^9^ and could be highly relevant to larger, complex blocks of genes identified in GWAS and eQTL studies.

Recent work has underscored that the genetic basis of PD is heavily rooted in the disruption of gene regulation within specific cell types. For instance, Dong et al. showed that PD risk variants are significantly enriched in non-coding regulatory elements, such as enhancers, that are active in dopamine neurons, pinpointing non-coding variation as a primary disease mechanism^16^. Building on this principle, a recent study by Lin et al. (manuscript submitted) systematically linked risk variants to their functional targets, identifying 124 high-confidence "eGenes" for PD. Critically, this work revealed that for most variants, the functional eGene was not the nearest physical gene, powerfully illustrating the complexity of *cis*-regulatory landscapes. However, these foundational studies have focused almost exclusively on SNPs and small indels, leaving the contribution of larger SVs to these regulatory circuits mostly unexplored. Recent evidence confirmed that SVs are a potent source of regulatory QTLs in bulk human brain tissue^8^. Preferential vulnerability of specific cell types however is a hallmark feature distinguishing human neurodegenerative diseases. For example, inclusions are preferentially found in glutamatergic and dopaminergic neurons of PD^17,18^, while inclusions preferentially affect glia cells in multiple system atrophy^19^. Thus, resolving their impact at a cell type resolution represents an important next step.

Here, we unite these technological and biological advances. By generating new long-read whole genome sequencing (WGS) data for the Parkinson’s Cell Atlas in 5D (PD5D) (Lin et al., manuscript submitted) cohort and integrating them with the existing single-nucleus RNA sequencing (snRNA-seq) dataset (Fig. 1a), we perform the first study, to our knowledge, for any disease, to link SVs to their effects on cell type-specific gene expression, using both eQTL and allele-specific expression (ASE) analysis, spanning health, disease onset, and progression. Our analysis uncovers candidate SV-gene associations missed by previous studies and prioritizes a set of functional SVs for future therapeutic investigation.

**Fig. 1.**
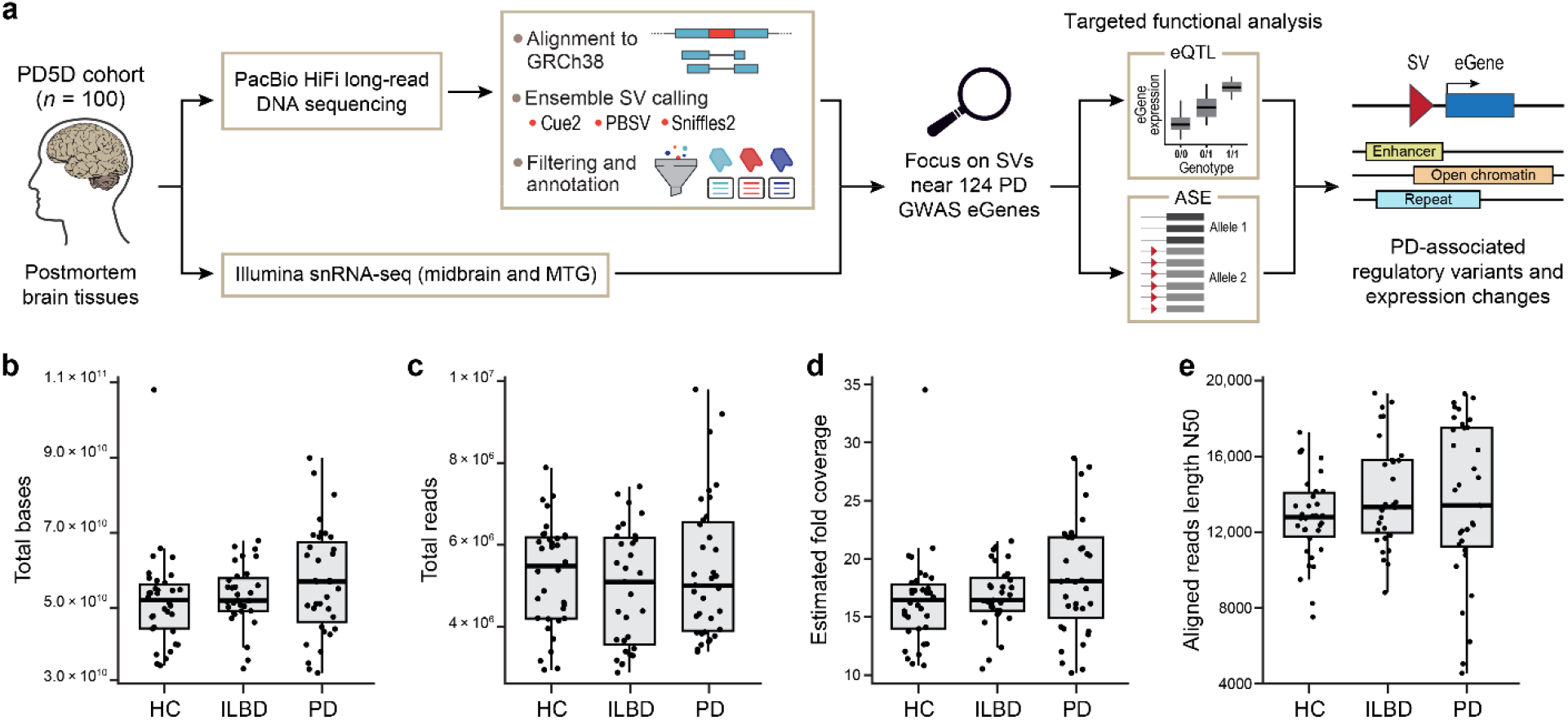
Study overview and sequencing metrics. **a**, Graphical overview of the study, showing flow of data from DNA samples to PacBio HiFi long reads followed by computational processing including alignment to GRCh38, ensemble variant calling using multiple callers, filtering and annotation, and targeted eQTL and ASE analyses integrating snRNA-seq data from the same samples. Our targeted approach focused on 124 causally associated PD GWAS "eGenes." The main output of this workflow is a list of SVs significantly associated with the expression of nearby PD eGenes. **b-e**, Box plots showing the quality of HiFi WGS data. Samples are grouped based on clinical diagnosis (x-axis). Data points show values from each sample. Sequencing read quality metrics plotted are indicated on the y-axis.

## Results

### Generating long-read sequence data for the PD5D cohort

To comprehensively characterize SVs impacting PD, we generated HiFi long-read WGS data from the PD5D cohort. DNA was isolated from brain tissue and sequenced using the PacBio Revio platform to produce HiFi long-read data (Methods). This final cohort consists of 100 individuals, including healthy controls (n=34), those with incidental Lewy body disease (ILBD) that could be considered prodromal PD (n=31), and confirmed PD patients (n=35) (Supplementary Table 1). These groups were similar in age and sex to prevent demographic factors from confounding the results.

Long-read sequencing of DNA from these PD samples generated a median of 52.63 Gb across 5.22 million reads per sample (Fig. 1b,c), corresponding to a median estimated fold coverage of 17x (Fig. 1d), and produced a deep and contiguous dataset. The median aligned read length N50 was 13.26 kb (Fig. 1e), underscoring the high contiguity of the sequencing data and ensuring confident alignment to the genome. Further evidence for the overall high quality of these data is shown by additional metrics (Extended Data Fig. 1), particularly the high alignment rates across all samples, with a median of 99.96% of reads successfully aligned to GRCh38 (Extended Data Fig. 1g; Supplementary Tables 2-3). Only three samples were removed for quality reasons (Methods; Supplementary Table 2). Collectively, these metrics demonstrate the generation of a high-quality, deep-coverage long-read sequencing dataset, which is a critical prerequisite for the SV calling pipeline.

### SV calling and a pipeline for generating a high-confidence SV callset

To generate a comprehensive and high-quality catalog of SVs for our cohort, we established and applied an evidence-driven ensemble variant calling and multi-stage processing pipeline (Fig. 1a and Extended Data Fig. 2a,b). Because no single SV caller has emerged yet as a “gold standard”^8,20^, we chose the following ensemble approach. For each of the 100 individuals, we generated initial SV calls using three distinct tools: Cue2 (a new release of Cue^21^ optimized for long reads), PBSV^22^, and Sniffles2^23^ (Methods; Extended Data Fig. 2a). The per-sample calls were first filtered to remove small variants and split by variant type (insertion, deletion, duplication, and inversion) to prevent merging of different types. We then sequentially combined variants first across individuals, then across callers (merging near-duplicate records), and finally across variant types using SURVIVOR^24^, yielding a unified list of 84,525 SVs.

### Ensemble SV calling strategy reveals caller-specific patterns

An initial analysis of the callsets revealed that the degree of agreement between the three SV callers varied substantially by variant type (Fig. 2a). Deletions showed the highest concordance (measured as the percentage of SVs commonly identified across callers out of all SVs identified by at least one caller, similar to the Jaccard Index), with a large proportion (58%) identified by all three callers. Insertions showed the next highest concordance between PBSV and Sniffles2 (54%). As Cue2 is not designed to report insertions, most insertions achieved the maximum possible support of two callers. Interestingly, we noticed that some insertion SVs (16%) identified by PBSV and Sniffles2 were matched with duplications identified by Cue2 (Methods). By contrast, concordance was substantially lower for duplications and inversions, underscoring the difficulty in detecting these event types. For both of these SV types, Cue2 identified more SVs than the other two callers combined.

**Fig. 2.**
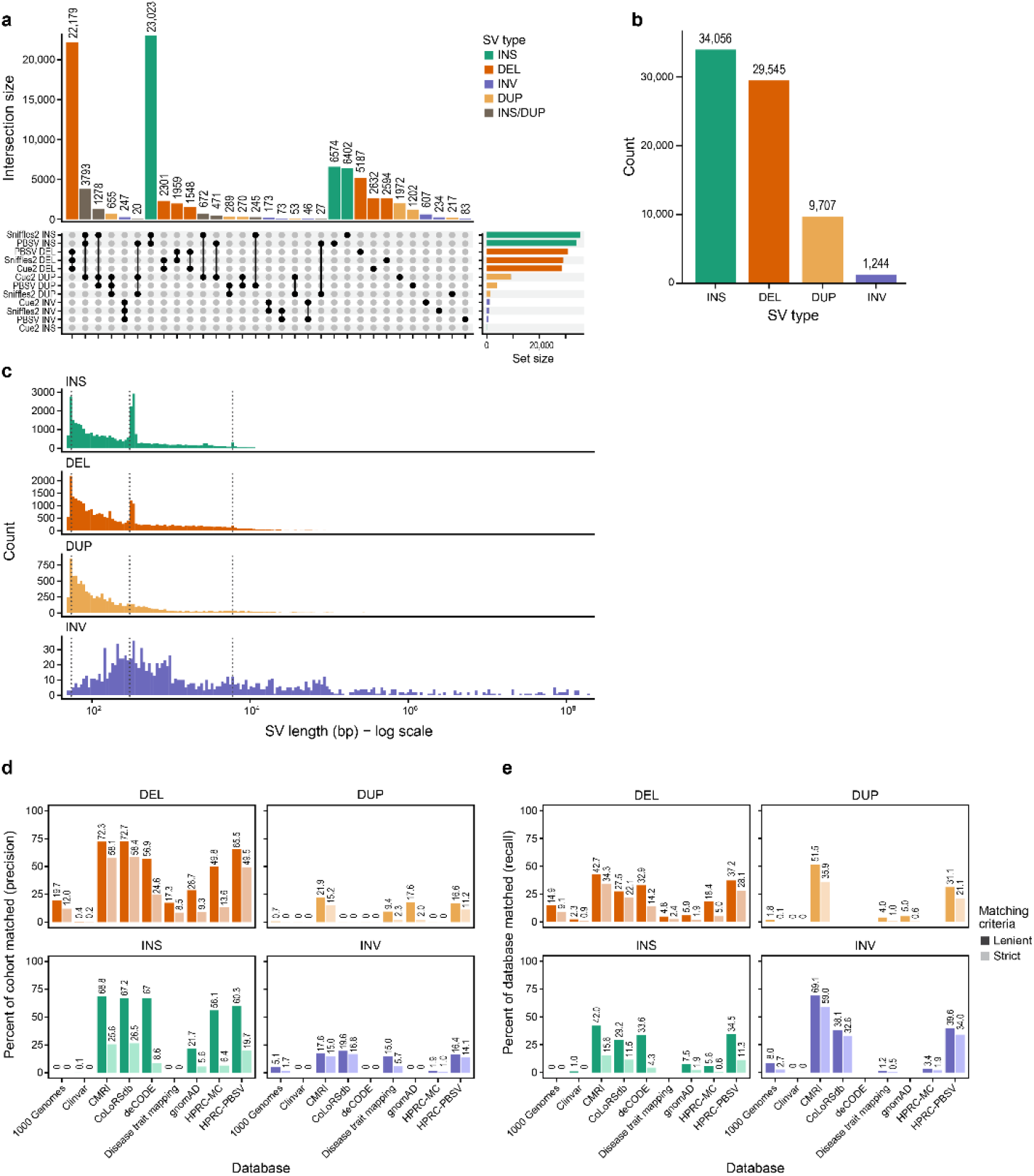
SV characterization. **a**, Upset plot showing the number of SVs commonly or uniquely called across the three callers for each SV type. SVs classified as an insertion by at least one caller and a duplication by at least one other caller are shown as INS/DUP here. The bottom right bubble plot shows set membership. Black circles indicate which sets are included in a particular intersection. Caller name and SV type are indicated in each row. Upper bar chart shows the size of each intersection. Set size bar chart (right) shows the total number of SVs in each individual set. Both bar plots are color-coded by SV type. **b**, Bar chart showing the composition of high-confidence SVs. SV types are color-coded and indicated on the x-axis. Number of SVs are on the y-axis, and the exact count of each type is specified on top of each bar. **c**, Histogram showing the distribution of SV sizes by SV type. SV type is color-coded and indicated on each subpanel. X-axis shows the size of SVs on a log_10_ scale. Y-axis shows the number of SVs. Grey dotted vertical lines are placed at 60, 300, and 6,000 bp in size to show SV lengths associated with replication slippage, Alu-, and LINE-mediated events, from smallest to largest, respectively. **d**,**e**, Bar charts showing precision, **d**, and recall, **d**, of SVs identified in this study against public SV catalogs. Each panel shows different SV types and public repositories are specified on the x-axis. In each comparison, precision shows the percentage of (SVs in the database matching SVs in this study) divided by (all the SVs in this study). Recall shows the percentage of (SVs in the database matching SVs in this study) divided by (all the SVs in the database).

This pattern of discordance likely reflects the fundamental differences in each caller’s algorithmic approach. For example, PBSV and Sniffles2 are alignment-based methods that extract SV signals from read alignments (e.g., split reads and read depth) and rely on heuristics to call SVs, which often fail to accurately model more complex SV signatures, especially in the presence of alignment artifacts that often impact duplications and inversions^25^. By contrast, Cue2 is a data-driven approach, which leverages deep learning to discover SVs in custom multi-channel images that encode SV-informative signals (including signals recovered from alignment artifacts), enabling it to handle more complex SV patterns in the data.

### Composition and size distribution of high-confidence SVs

The initial callset of 84,525 SVs was subsequently refined by removing 6,768 (8%) SVs located within challenging genomic regions known to be prone to false-positive calls^26^, namely segmental duplications (repeats > 1kb), assembly gaps, and centromeres (Methods; Extended Data Fig. 2b). After further removing 3,205 low quality SVs based on read-based metrics (Methods; Supplementary Table 4), the final callset contained 74,552 high-confidence SVs. Our final, high-confidence callset is predominately insertions (34,056) and deletions (29,545), which together account for over 85% of all detected SVs (Fig. 2b). Duplications (9,707) and inversions (1,244) were substantially less frequent. This observed distribution, where insertions and deletions constitute the majority of SVs, is highly consistent with SV callsets from large-scale studies^1,8,9,27^. The size distribution analysis revealed distinct, mechanism-driven patterns for each SV type as observed in other studies^8,10^ (Fig. 2c). Both deletions and insertions have noticeable peaks at about 50 and 300 bp. This pattern is in line with the expected SV size distribution from biological events. Specifically, replication slippage likely results in small (50-100 bp) SVs^1^ and Alu element differences explain a smaller ∼300 bp peak^28^. An even smaller peak at ∼6000 bp is evidence of long interspersed nuclear element (LINE) differences^29^ (Fig. 2c). Duplications also showed a high frequency of small events (<100 bp) similar to insertions, suggesting these small events might share common mutational mechanisms. By contrast, the inversions were more broadly distributed across sizes. The recapitulation of these distinct, mechanism-driven size distributions lends strong confidence to our dataset’s quality and biological accuracy.

### Region-based annotation of identified SVs

Given the breakpoints of identified SVs, we used genome annotation information to characterize SVs’ potential to affect gene expression. We first looked at the genome locations where SVs are most often found (Extended Data Fig. 3a). Cytobands identified as most SV-prone are strongly consistent with established genomic “hotspots” known for inherent instability^30^. Many of these regions are known fragile sites and have been implicated in diseases. For example, the top two cytobands with the most SVs, 19p13.3 and 5p15.33, are associated with a microdeletion/microduplication syndrome^31^ and cross-cancer risk^32^, respectively. Next, to identify SVs with potential functional impact, we compared the positions of our SVs with known regulatory regions in functional genomics databases using AnnotSV^33^ (Methods and Supplementary Table 5). Importantly, to specifically identify brain-active regulatory elements, we curated GeneHancer^34^ data using tissue expression to select for regulatory elements expressed in brain (Methods). This identified 6,225 (8.35% with 9.64% expected overlap with random SV genomic intervals; Methods) SVs overlapping regulatory elements and 1,036 (1.39%) SVs overlapping brain-active promoters and enhancers (Extended Data Fig. 3b,c). We extended the overlap analysis to other functionally relevant regions annotated for GRCh38, including repeat regions from RepeatMasker (62.7%; 61.5% expected at random), transcription factor binding sites (36.9%; 36.2% expected at random), histone methylation (63.4%; 77.4% expected at random), and acetylation (70.3%; 86.0% expected at random), CpG islands (5.0%; 1.7% expected at random), DNase hypersensitivity (20.1%; 27.5% expected at random), and evolutionary conserved regions (91.9%; 98.7% expected at random) (Methods, Extended Data Fig. 3c). The nearly three-fold enrichment of SVs at CpG islands relative to random expectation suggests SVs may have an impact on regulatory and promoter elements. Overall, the extensive overlap with evolutionarily conserved regions and active chromatin marks suggests potential functional consequences for these SVs and indicates that some affect biologically important genomic regions.

### Comparing SVs identified in this cohort to established databases

To identify novel SVs discovered in this study, we queried nine SV-containing databases using Truvari^35^ (Methods; Fig. 2d,e). Our SVs showed greater concordance with databases that leveraged long-read sequencing, namely the Human Pangenome Reference Consortium (HPRC), Children’s Mercy Research Institute (CMRI), CoLoRSdb, and deCODE, compared to databases predominantly derived from short-read data like the 1000 Genomes Project and gnomAD (Supplementary Table 6). When checking whether two SVs are actually the same, the current best practice is to use inexact breakpoints match and consider sequence similarity^35,36^. Despite base-level precision of many callers, it is often difficult to resolve the exact position of breakpoints. For example, due to the repeated nature of tandem repeat sequences, callers may assign deletion breakpoints at different locations using the same evidence.

To test the impact of match stringency, we compared two methods: strict criteria that requires near-perfect breakpoints and sequence match, and lenient criteria that is more standard (Methods). By comparing precision and recall for each database using these criteria (Fig. 2d,e), we found that strict method captures substantially less matching SVs (precision was 26.0% for deletions and 10.3% for insertions; mean for the nine databases) compared to the lenient method (precision was 42.6% for deletions and 37.9% for insertions; mean for the nine databases). We observed a marked difference in the match rate by SV type. By contrast with insertions and deletions, duplications and inversions showed consistently lower match rates across all databases. This discrepancy could be explained by the lack of uniformity in how SV types are classified and reported across the various reference databases (Extended Data Fig. 3d), complicating the process of matching SVs regardless of comparison parameters.

### Selection of SVs for functional analysis

To conduct a robust functional analysis of SVs in PD, it is essential to first generate highly accurate genotype calls. In an ensemble calling strategy, the accuracy of assigned genotypes may vary among the callers. We first assessed the degree to which genotypes disagreed on a subset SVs identified by more than one caller across SV types (Supplementary Fig. 1a). This revealed that for all SV types, the genotype agreed for about 90% of SVs called by both callers (pairwise comparisons), underlining the importance of assigning accurate genotypes for the ∼10% of SVs with genotype discordance. We benchmarked the performance of SV genotypers Kanpig^37^ and VaPoR^38^ using synthetic datasets and used this information to determine our genotype verification strategy (Methods; Supplementary Note 1). The advantages of this variant calling and genotyping approach are: (1) we leverage the distinct advantages of three variant callers to establish a single cohort-level VCF; and (2) we consistently assign confident genotypes using Kanpig^37^, which decouples genotyping from discovery to rigorously assess allelic presence using variant graphs and k-mer vectors. This method is able to resolve complex loci and neighboring SVs that frequently confound variant callers^37^.

Generally, the large number of SVs, observed in the range of tens of thousands per individual, may lead to missing many associations of potential interest due to multiple hypothesis corrections in functional analyses^1,3^. Therefore, we elected to use a targeted strategy to focus the analysis on SVs located in the vicinity of PD GWAS loci, narrowing our analysis to SVs within 100 kb of 124 functionally identified high-confidence PD “eGenes” identified by previous SNP-based analyses (Lin et al., manuscript submitted). We elected this strategy because we wanted to assess whether SVs are potentially responsible for these gene-PD associations. This approach is also motivated by the insight that PD risk variants are significantly enriched in non-coding regulatory elements, such as active enhancers, suggesting the possibility that the effect of these variants on PD risk is due to how they regulate expression of genes of interest^16^.

SV allele frequency (AF) and cohort size also affect statistical power. SVs must be sufficiently frequent in a cohort for reliable statistics. Especially in a small cohort like the PD5D cohort included in this study (n=100), rare variants (defined here as AF < 1%) are not sufficiently prevalent for functional analysis. After identifying 727 SVs within 100 kb of 84 PD GWAS eGenes (Fig. 3a; 40 eGenes did not have any SVs within the 100 kb window), we removed 323 SVs with valid genotype calls in less than 5 samples, less than two copies of the alternate allele, or present only in healthy control individuals (Fig. 3b; Supplementary Table 7). Our selection strategy yielded 404 SVs for functional analysis (218 insertions, 176 deletions and 10 inversions; Fig. 3c). Thirty three of these 218 insertions were identified as insertions by PBSV or Sniffles2, but could be duplications as Cue2 identified a duplication in the same location (Supplementary Table 7).

**Fig. 3.**
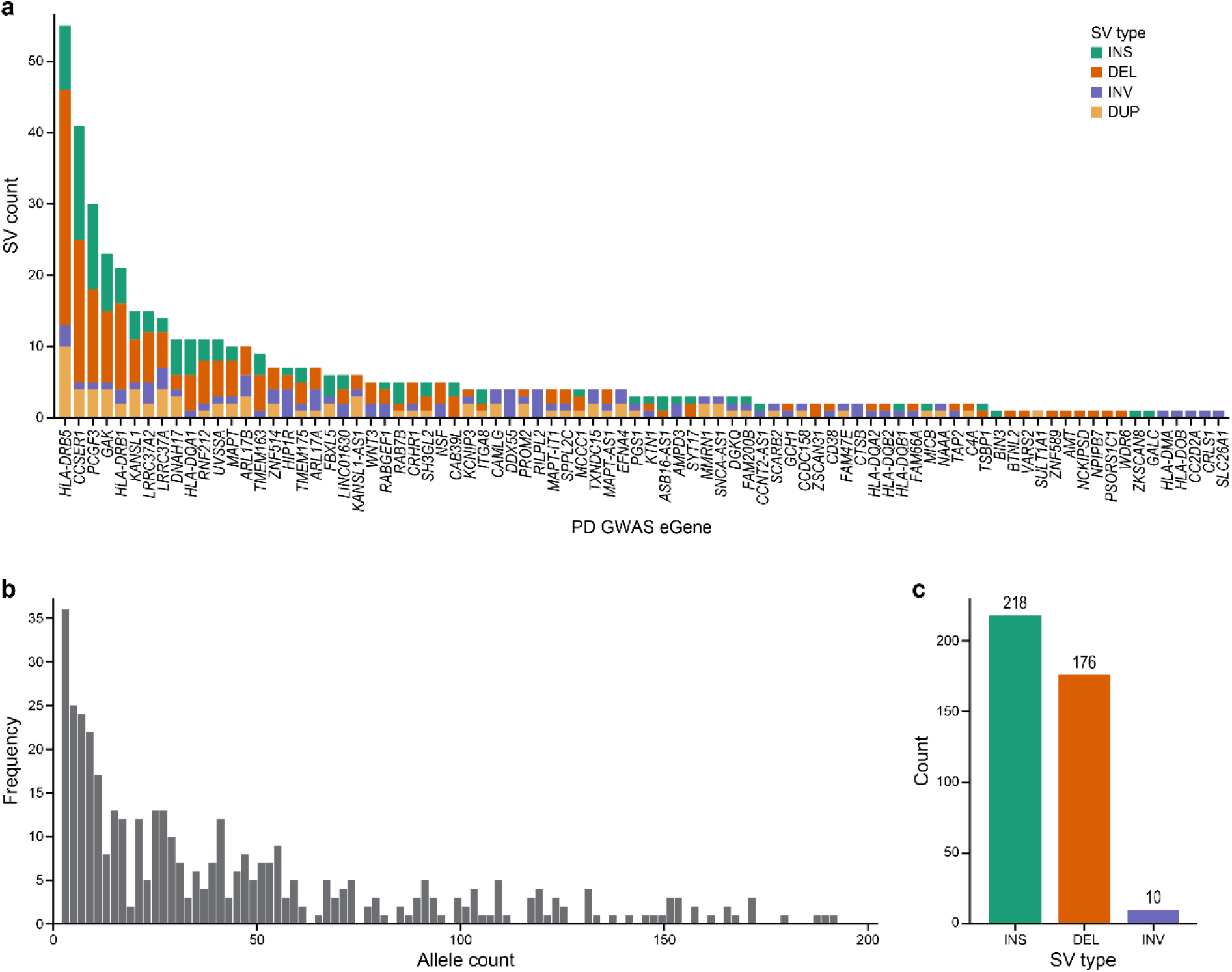
PD eGene SV characterization. **a**, Bar chart showing the number of SVs within 100 kb of PD GWAS eGenes. Y-axis shows the SV count. Bars are color-coded by SV type. **b**, Histogram showing the frequency of alternate allele in this study. X-axis shows the alternate allele count and y-axis shows the frequency. **c**, Bar chart showing the composition of SVs passing QC. SV types are color-coded and indicated on the x-axis. Number of SVs are on the y-axis and the exact count of each type is specified on top of each bar.

### Structural variants identify eQTLs in specific cell types

To investigate the functional impact of these SVs on gene regulation in the brain, we performed expression quantitative trait loci (SV-eQTL) analysis, correlating SV genotypes with gene expression levels in nine and eleven major cell types of the middle temporal gyrus (MTG) and midbrain, respectively. We used covariance (sequencing batch, age, sex, PMI, RIN, and hidden relatedness among individuals) adjusted pseudobulk expression from snRNA-seq data of the matched individuals (Methods).

Our analysis identified 173 unique SVs (286 SV-gene pairs) with significant *cis*-regulatory effects on gene expression in MTG and midbrain across all cell types assessed, many of which appear specific to a single cell type (Fig. 4, Extended Data Fig. 4a, 5, Supplementary Table 8). An alternative approach to finding eQTLs explicitly correcting for population stratification is presented in Supplementary Note 2. Among these SV-gene pairs, 43.0% (123 pairs) showed significant effects in both brain regions, and 67 and 96 pairs were uniquely significant in MTG and midbrain, respectively (Extended Data Fig. 5a,b). In both regions, SV eQTL results are dominated by the same three genomic regions. First, the 17q21.31 region has SV eQTLs with strong effects in multiple cell type signals affecting the putative functional risk genes for PD nominated in this locus^16^ (Lin et al., manuscript submitted), including *ARL17B*, *KANSL1* and *LRRC37A*. Previous fine-mapping and functional studies have established the 17q21.31 H1 haplotype as a significant risk factor for PD, implicating *KANSL1* in autophagy and lysosomal dysfunction and *LRRC37A* in astrocytic clearance of alpha synuclein, suggesting these genes drive disease susceptibility alongside *MAPT*^16,39–41^. Our results show that the same highly significant SVs, including insertions (e.g., chr17_46252361_INS_828), deletions (e.g., chr17_46237501_DEL_-724), and the 45,106 bp inversion (chr17_46231903_INV_45106), are associated with expression of these genes in both brain regions (Fig. 5a, Extended Data Figs. 6a,b and 7a,b). Second, the Human Leukocyte Antigen (HLA) region on chromosome 6 has the most *cis*-eQTLs (87 and 77 significant SV-gene pairs across 48 and 51 unique SVs in MTG and midbrain, respectively) (Extended Data Fig. 4a). While we observed numerous HLA genes (including genes encoding both HLA-DR and HLA-DQ protein families) associated with SVs in this locus, this region’s notoriously complex linkage disequilibrium and haplotype structures limit our confidence in the identification of causal variants without additional investigation. Third, the *CRIPAK* and *UVSSA* locus on chromosome 4 has multiple SVs associated with expression of these genes (e.g., chr4_1372768_INS_323 and chr4_1324644_DEL_-94) in both brain regions (Fig. 4, Extended Data Figs. 6c, and 7c,d). However, the cell types in which significant effects were detected differed by the region. Interestingly, while SVs are uniformly associated with up-regulation of these genes in glutamatergic and GABAergic neurons in the MTG, the midbrain results showed association with both up- and down-regulation of these genes in microglia and oligodendrocytes. The observed down-regulation was specific to *UVSSA* in midbrain (Extended Data Figs. 6d and 7c-e).

**Fig. 4.**
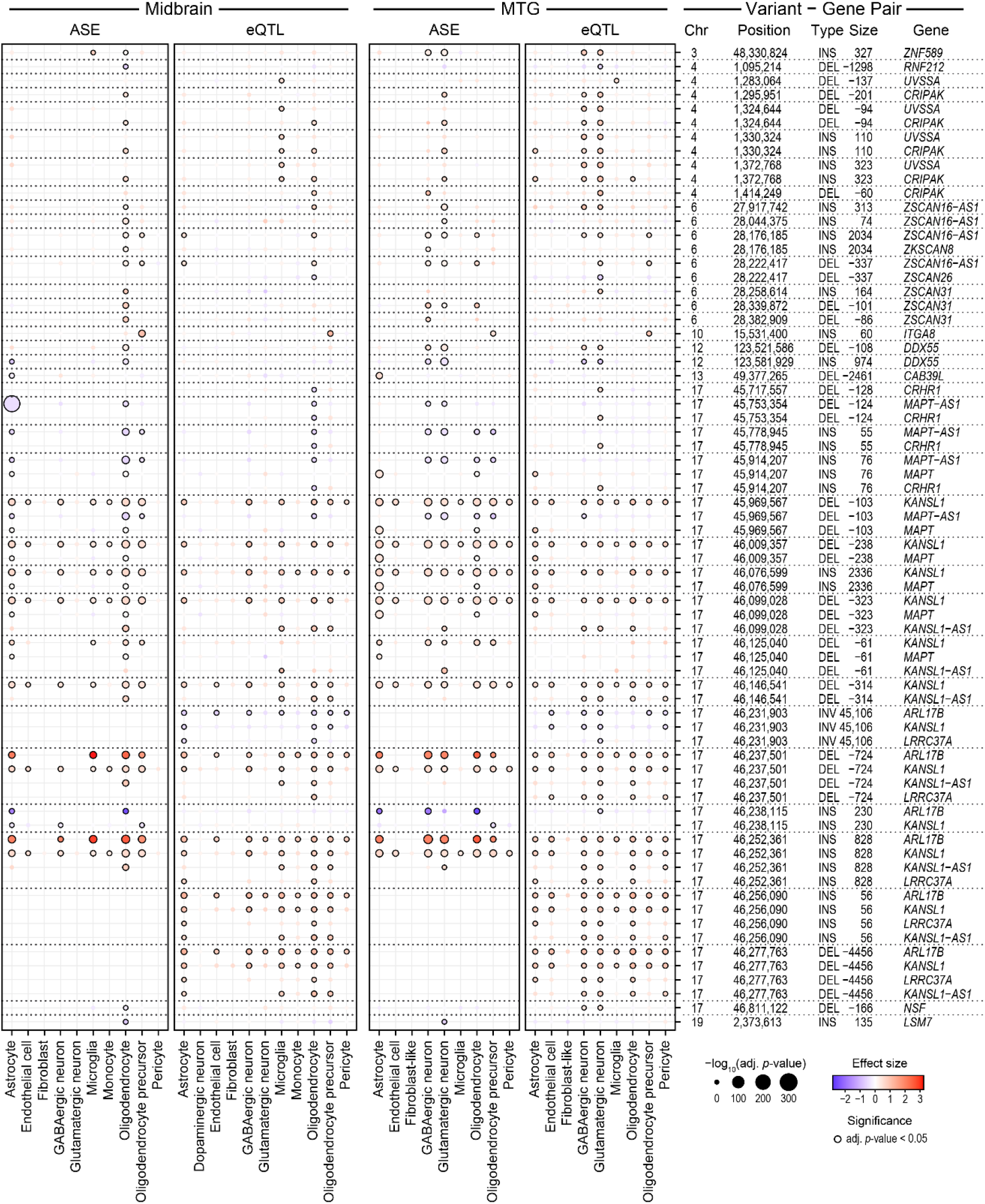
ASE and eQTL analysis of SVs near PD eGenes. **a**, Bubble plots showing ASE and eQTL analyses in midbrain (left) and MTG (right). Only significant SV-gene pairs in both regions are plotted. SVs in the HLA locus (chr6: 28,510,120-33,480,577) are not shown for clarity. Information about SV-gene pairs is summarized in row names in a 5-column format (chromosome, position, SV type, SV length, and gene name). Cell types tested are shown on the x-axis. Adjusted p-value is shown by the bubble size (-log_10_ scale) and effect size is shown by the color. Significant associations have a black outline. Same SV affecting multiple genes is shown by the dotted horizontal lines. SV-gene pairs on the y-axis are sorted by chromosome and position.

**Fig. 5.**
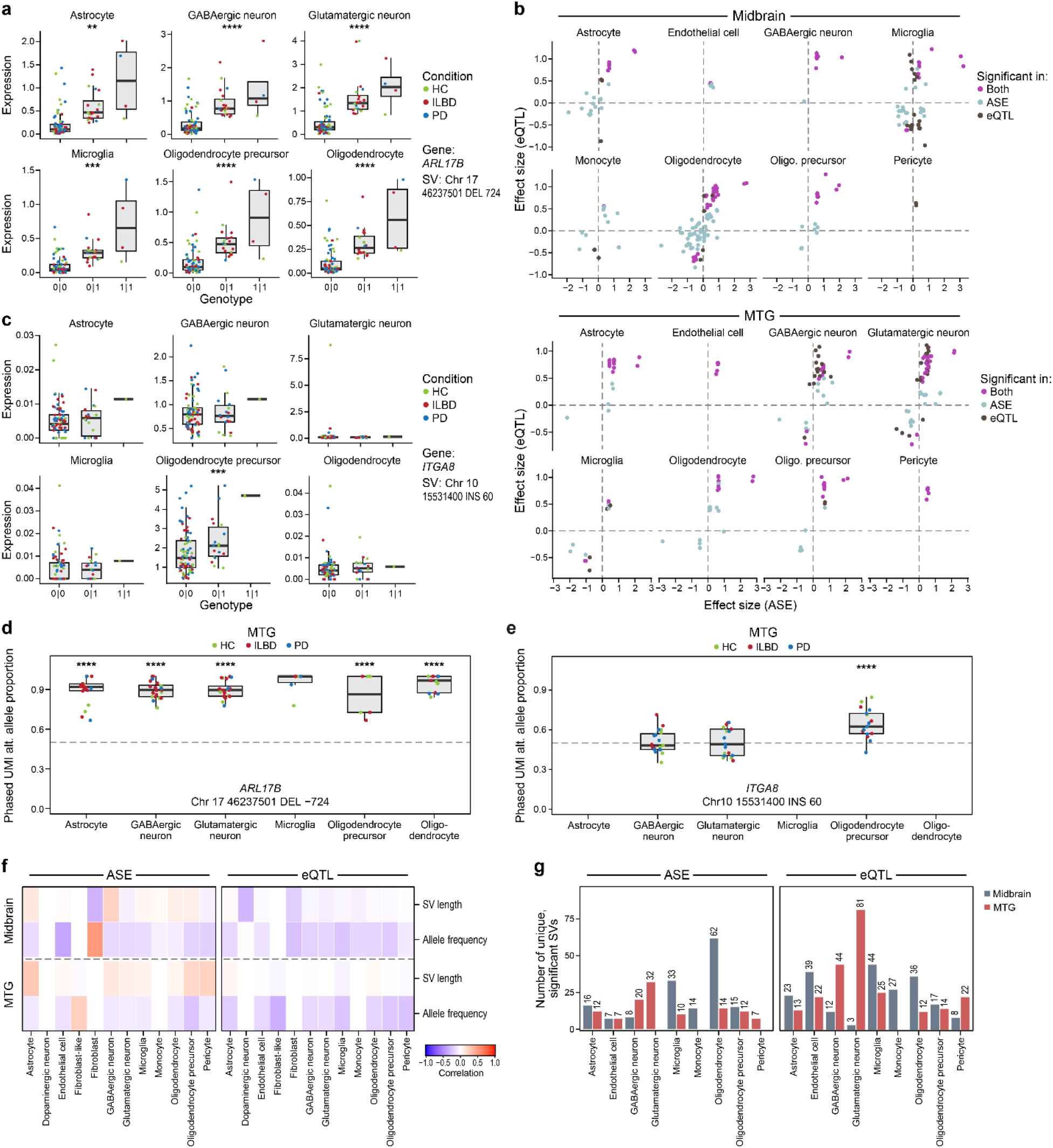
ASE and eQTL results comparison. **a**,**c**, Box plots showing the pseudobulk expression of SV-eQTL significant target genes from snRNA-seq data across cell types in MTG. SV-gene pair and cell types are indicated on the right. Individuals are grouped into SV genotype (x-axis) and their averaged expression values (y-axis) are plotted. Individual data points are color-coded by the diagnosis. **a**, an example of a shared cell type effect. **c,** an example of a specific cell type effect. **b**, Scatter plots comparing the effect sizes from ASE and eQTL analyses across different cell types in midbrain (top) and MTG (bottom). Effect sizes from ASE are shown on the x-axis, and those from eQTL are shown on the y-axis. Dots are color-coded by significance in the two methods. Only SV-gene pairs significant in at least one method are plotted. **d**,**e**, Box plots showing ASE in MTG. X-axis shows proportion of phased UMIs mapping to alternate allele. Y-axis shows cell types. Individual data points are color-coded by the diagnosis. **d**, an example of a shared cell type effect. **e,** an example of a specific cell type effect. **f**, Heatmaps showing the correlation between ASE or eQTL effect size and SV properties across cell types in midbrain (top) and MTG (bottom). The test that generated the effect size is indicated at the top. SV properties include SV length and minor allele frequency (y-axis). Cell types are shown on the x-axis. **g**, Bar charts showing number of unique significant SVs from ASE (left) and eQTL (right) analyses in the midbrain (black) and MTG (red). The number of significant SVs is on the x-axis and cell types are listed on the y-axis. Statistical significance in **a**, **c**-**e** was indicated by asterisks based on adjusted p-value thresholds: * Padj <= 0.05, ** Padj <= 0.01, *** Padj <= 0.001, and **** Padj <= 0.0001. No asterisk means Padj > 0.05.

Likewise, while some PD risk loci, such as those within the HLA region and the *MAPT*/*KANSL1* locus, showed regulatory activity in both the midbrain and MTG, many specific SV-gene associations were detected in only one of the two regions. In the midbrain analysis, SV eQTLs absent or less prominent in the MTG results (Extended Data Fig. 5a) included genes such as *SCARB2* (associated with chr4_76150170_DEL_-109). While *SCARB2* was expressed in both regions (Supplementary Fig. 2a,b), its association with the SV was only significant in the midbrain (Extended Data Figs. 6e and 7f). More specifically, *SCARB2* was significantly up-regulated in oligodendrocytes in the midbrain (midbrain eQTL padj = 0.0012 with effect size = 0.78), but not in MTG (eQTL padj = 0.15). By contrast, several SV-eQTLs were found to be significant only in the MTG (Extended Data Fig. 5b), such as *KTN1* (associated with chr14_55489789_DEL_-65) (Extended Data Figs. 6fand 7g). Similar to the *SCARB2* example, the expression of *KTN1* alone does not explain the difference between the two brain regions (Supplementary Fig. 2a,b). These findings indicate that while many SV-eQTLs located near PD eGenes are functionally relevant in both the MTG and midbrain, the specific gene targets and regulatory vulnerability may differ between these two brain regions, emphasizing the necessity of performing this analysis in multiple tissue contexts. While it is possible that these findings are due to regulatory differences, other reasons such as a lack of power due to a low frequency of cells in some cell types in one brain region are also possible.

Another interesting association was observed for two SVs (chr12_123521586_DEL_-108 and chr12_123581929_INS_974) and *DDX55*. While both SVs were significant eQTLs in the MTG, the deletion was associated with increased expression and the insertion with decreased expression in all cell types significantly affected (Extended Data Fig. 7h,i). In 16 samples for which one copy of both SVs was detected and phased, the SVs were always present in trans rather than in cis at the DDX55 locus. Specifically, the insertion was associated with significant down-regulation of *DDX55* in glutamatergic and GABAergic neurons in the MTG (eQTL effect size = -0.72 and -0.72; padj = 5.36e-03 and 9.49e-03, respectively). This insertion is an LTR5Hs element of the HERV-K endogenous retrovirus (Methods; Dfam bit score = 1215.9 with E-value = 0) and it was inserted into a AluSx1 repeat element present ∼20 kb upstream of *DDX55* (Extended Data Fig. 10a). Without additional experiments, it is not possible to determine a mechanism for the association of these SVs with *DDX55* expression, but there are potential clues in the existing annotation of this genomic region. This insertion is in a region annotated as having weak enhancer activity based on its chromatin state segmentation by hidden Markov model (HMM) from data of nine human cell lines^42^. Also, this insertion is within 100 bp of open chromatin regions across multiple cell types in dorsolateral prefrontal cortex from snATAC-seq data in brainSCOPE^43^. It is possible, though speculative, that the insertion disrupts enhancers located in one of these regions to decrease DDX55 expression. By contrast, the deletion was significantly associated with up-regulation of *DDX55* in the same neuronal populations (eQTL effect size = 0.60 and 0.66 with padj = 2.00e-02 and 1.56e-02 in glutamatergic and GABAergic neurons, respectively). This deletion removes a simple repetitive DNA sequence found in the 3’ end of long interspersed nuclear element-1 (LINE-1) (Extended Data Fig. 10a). Although there is a brain-active enhancer and open chromatin regions nearby, it remains unclear whether this deletion exerts its effect through the modulation of these elements or via other mechanisms.

### Structural variants identify ASE differences in specific cell types

We complemented our eQTL analysis with ASE analysis to identify allelic imbalance in individuals heterozygous for the SVs. This method offers increased statistical power due to its approach, which compares the expression of maternal and paternal alleles within a single individual. ASE analysis avoids many sources of inter-individual variability and confounding that affect eQTL comparisons^12,44,45^. The ASE results identified mostly the same SV-gene pairs outside of the HLA locus compared to the eQTL results, recapitulating the strong effects on gene expression in multiple cell types at 17q21.31 and multiple SVs significantly associated with *CRIPAK* and *UVSSA* genes on chromosome 4 (Fig. 4, Extended Data Fig. 7a-e, Supplementary Table 9a,b). By contrast, in the HLA locus, especially in the MTG, the two methods largely disagreed (Extended Data Fig. 4a), which could be due to difficulties in phasing the HLA region, lower expression levels for these genes in some cell types decreasing power for ASE analysis (Extended Data Fig. 4b,c), or other unidentified issues. Next, we directly compared the effect sizes from the eQTL and ASE analyses in the MTG and midbrain (Fig. 5b). This revealed a high agreement between the two methods, with a strong positive correlation in effect sizes for SVs identified as significant by both approaches (Pearson correlation r = 0.62 and 0.65 across all cell types in MTG and midbrain, respectively). Those SVs significant in only one of the two tests showed mostly consistent fold change direction, increasing our confidence in these regulatory signals. In MTG, the effect size correlation was stronger in SVs found significant by only one method compared to those found by both methods (Pearson correlation r = 0.65 and 0.79 among SVs significant only in ASE or eQTL, respectively). In contrast to the MTG, this effect size correlation of SVs found significant by only one method in midbrain showed less agreement (Pearson correlation r = 0.47 and -0.02 in ASE or eQTL, respectively).

While ASE and eQTL results exhibited high concordance in the direction of effects, they often did not agree on the significance. Such divergence underscores how these methods report distinct aspects of genetic regulation. While ASE could avoid population-level biases, the analysis is inherently restricted to the subset of individuals heterozygous for both the SV and the informative SNPs. By contrast, eQTL analysis utilizes the entire cohort, which may provide more power when the number of heterozygous individuals is limited. Furthermore, ASE requires sufficient read depth over heterozygous SNPs to calculate an allelic ratio. Even for highly expressed genes, if there is a lack of strong linkage between the SV and the informative SNPs used for counting, the ASE signal may be attenuated compared to the total expression change detected by eQTL. It is also possible that issues with the phasing of the SVs could lead to disagreement between ASE and eQTL analysis. Other factors may also contribute to divergence between eQTL and ASE analyses. We highlight two types of examples of this divergence. First, a major distinction lies in the regulation of *MAPT-AS1*. While several SVs in 17q21.31 (e.g., chr17_45778945_INS_55 and chr17_45969567_DEL_−103) down-regulated *MAPT-AS1* expression in multiple cell types including both glial and neuronal types in the MTG, their effects were only identified to be significant with ASE (Extended Data Figs. 6g,h and 7). The eQTL analysis did not detect significant changes in the MTG on *MAPT-AS1*. Similarly, in the midbrain, while the ASE method identified significant down-regulation of *MAPT-AS1* in oligodendrocytes, OPCs and astrocytes, the eQTL method only detected a significant effect in oligodendrocytes (Extended Data Fig. 6i,j). Second, certain SV-gene pairs were only detected as significant by the eQTL method. Specifically, *ARL17B, KANSL1, LRRC37A, and KANSL1-AS1* were found to be regulated by chr17_46231903_INV_45106, chr17_46256090_INS_56, and chr17_46277763_DEL_-4456 (Fig. 4). Notably, while these genes show significant allelic imbalance with other SVs, the specific regulatory influence of these three SVs on these target genes did not reach significance in our ASE analysis. These differences could be due to (1) remaining technical confounders not being corrected for in the eQTL analysis, (2) reference bias in the ASE analysis, (3) biological events that lead to the imbalance in alleles’ expression without significant change in the gene’s total expression, and (4) ASE and eQTL measurements derived from different gene isoforms. These findings emphasize the value of performing both types of analyses.

To overcome potential snRNA-seq sequencing depth limitations contributing to such discordance, we generated a targeted MTG and midbrain snRNA-seq dataset using hybrid selection^12^ (Methods) to specifically enrich for reads mapping to a subset of our prioritized PD risk loci and performed ASE analysis (Extended Data Fig. 8a,b). Hybrid selection targeted an overlapping, but not identical set of genes for each brain region (Supplementary Table 10). This targeted ASE approach revealed mostly consistent effect sizes compared to ASE using non-selected snRNA data (Pearson correlation r = 0.958 and 0.933 in midbrain and MTG among SV-gene pairs that are significant in at least one of the two methods, respectively; Extended Data Fig. 9a,b). For targeted genes in MTG and midbrain, we observed more significant SV-gene pairs with hybrid selection than without hybrid selection (291 vs. 113 and 180 vs. 154, respectively (Supplementary Table 9 and Extended Data Fig. 9c)). As expected for genes not targeted by hybrid selection, we did not see enrichment in significant SV-gene pairs in MTG (29 with selection vs. 31 without selection) nor in midbrain (0 vs. 36). This finding confirms that hybrid selection can increase power to detect allelic imbalance.

### Comparisons of SV eQTL and ASE results

To illustrate the diverse regulatory patterns identified in this study, we highlight several examples (Fig. 5a,c-e, Extended Data Figs. 6, 7) in the context of existing genome annotations. For instance, a 724 bp deletion in 17q21.31 (chr17_46237501_DEL_-724) was associated with a significant increase in the expression of the nearby genes in both brain regions across multiple cell types, including both glial and neuronal lineages. Significantly affected genes include *ARL17B*, *KANSL1*, *KANSL1-AS1* and *LRRC37A*. This SV was previously reported in all of the long-read SV catalogs compared and in a disease trait-mapping study^46^, and was predicted to be benign by AnnotSV (Methods). In proximity to this deletion, there is a larger (45,106 bp) inversion (chr17_46231903_INV_45106), significantly affecting the same genes in multiple cell types (Fig. 4, Extended Data Figs. 6a,k and 7a,b). Despite its presence in CoLoRS and CMRI, there was not enough evidence to predict clinical significance for this SV. Our functional analysis showing its role as a strong eQTL in multiple cell types underscores its potential as a novel regulatory element and suggests it could be a good candidate for future mechanistic validation studies.

Next, we explored the relationship between SV properties and eQTL effect sizes across all cell types for each SV type. We tested two continuous variables related to SV: minor allele frequency (MAF) and SV length. Across all SV types combined (Fig. 5f), we observed a consistent negative correlation (ranging from -0.34 to -0.15 in MTG and - 0.27 to -0.089 in midbrain; except for dopaminergic neurons in midbrain with 0.013) between MAF and eQTL absolute effect sizes across all tested cell types. The modest effect we observe is consistent with a previous study that found low-frequency SVs (MAF 2–5%) had significantly larger effect sizes than SVs with higher MAFs in their dataset^9^. Furthermore, SV type-specific correlation analysis (Extended Data Fig. 9d) revealed distinct characteristics. For insertions, SV length showed weaker correlation with eQTL effect size compared to that of MAF across all cell types except for dopaminergic neurons (mean magnitude of Pearson r = 0.19 and 0.03 for MAF and SV length, respectively; Extended Data Fig. 9d). By contrast, deletions showed distinct patterns by the SV property. A consistent positive correlation was observed between SV length and eQTL effect size in 8 out of 9 cell types in MTG (Pearson r ranges from 0.036 to 0.32), and 10 out of 11 cell types in midbrain (Pearson r ranges from 0.16 to 0.446) (Extended Data Fig. 9d). Finally, for inversions, SV length and MAF showed opposite trends. SV length showed predominantly negative relationships across most cell types in the two regions (except in MTG astrocytes and oligodendrocytes). By contrast, MAF showed mostly positive correlation except in those two MTG cell types (Extended Data Fig. 9d). However, due to the small number of inversions passing our selection criteria, this correlation may not generalize for all inversions.

To assess the differential impact of SVs across specific brain cell types, we quantified the number of significant SVs detected within each cell type (Fig. 5g). When differences are observed between cell types, there are several possible reasons. First, the gene may not be expressed in all cell types. Second, there may be less power to detect SV effects due to lower numbers of cells for some cell types. Third, and most interesting, the SV may affect gene expression more in some cell types. To distinguish among these possibilities, we categorized specific effects based on gene expression and cell count thresholds (Methods; Extended Data Fig. 9e,f). Across both brain regions, there were 94 and 176 SV-gene pairs significant in only one cell type, from ASE and eQTL analyses, respectively. For 14 (14.9%) ASE and 26 (14.8%) eQTL significant SV-gene pairs, cell type specificity seems to be due to the gene’s expression being limited to the cell type showing the significant effect because (1) the expression levels are too low in other cell types and (2) there are enough cells in some of the other cell types. Additionally, for another 34 (36.1%) ASE and 72 (40.9%) eQTL SV-gene pairs, cell type specificity seems to be due to both limited expression in other cell types and not enough cells in those other cell types. The remaining 46 (48.9%) ASE and 78 (44.3%) eQTL SV-gene pairs are candidates for cell type specific SV regulatory effects. Examples of such candidates include a 60 bp insertion on chromosome 10 (chr10_15531400_INS_60) that was linked to a significant increase in *ITGA8* expression only in OPCs (eQTL FDR = 2.39e-07, ASE FDR = 8.68e-19, Fig. 5c,e, Extended Data Fig. 7k), and a 330 bp insertion on chromosome 8 (chr8_22625880_INS_330) that was associated with up-regulation of *BIN3* (Extended Data Fig. 7l). Notably, a recent study reported this SV near *BIN3* as a methylation QTL (chr8: 22,625,890; 329 bp intronic Alu element insertion)^9^. The coordinates and size of this SV are virtually identical to the SV that we identified as a significant eQTL specific to glutamatergic neurons (eQTL padj = 1.22e-02, ASE with selection padj = 9.74e-03) in MTG (Extended Data Figs. 5b, 7l and 8b). This insertion overlaps a chromatin accessibility peak identified in glutamatergic neurons that was not found in other cell types (Extended Data Fig. 10b). Previously, this SV was known to act in brain, but the relevant cell types were not known. Our finding of significant *BIN3* up-regulation specifically in glutamatergic neurons is a good example of how snRNA-seq provides insight into the relevant cell type for a variant.

Together, these findings underscore that integrating SV detection via long-read genomic sequencing with cell-type-specific functional assays is essential for comprehensively deciphering the complex genetic architecture underlying neurodegenerative diseases.

## Discussion

We applied HiFi long-read sequencing and cell type-resolved expression analysis to generate a high-confidence catalog of SVs and identify SV-eQTLs and SV associated allelic imbalance, revealing the important and highly specific role of SVs in gene regulation. To ensure that our SV catalog does not suffer from tool-specific blind spots and biases^47^, we designed an ensemble strategy that leverages the complementary strengths of multiple SV callers and subjects the consolidated callset to rigorous filtering and functional annotation, maximizing sensitivity without compromising the specificity required for downstream analysis. Our analysis suggests that SVs are a potent source of *cis*-regulatory variation and highlights their potentially cell type specific effects for PD related genes. From the 90 PD GWAS peaks, we focused on 124 eGenes and identified 173 SVs having significant associations with the expression of 51 and 38 of those genes in eQTL and ASE analyses, respectively. Although neuronal populations in the MTG, particularly glutamatergic neurons, have more significant SV-gene pairs (81 SVs), while glial cell types showed fewer associations (12-25 SVs, Fig. 5g), this may be due, in part, to gene expression level and cell number differences among cell types (Supplementary Fig. 2a-d, Extended Data Fig. 9e,f), or the genes selected for analysis (since they are based on PD GWAS loci). The eQTL analysis of midbrain endothelial cells highlights them as a potentially abundant source of significant SV-gene pairs with cell type specificity (Extended Data Fig. 9f). Furthermore, the investigation into SV properties showed a strong negative correlation between MAF and eQTL effect size in all but one cell type analyzed (Fig. 5f). Interestingly, we provided cell-type resolution for an SV near *BIN3*, a known PD locus, finding that the ∼330 bp insertion acts as a significant eQTL with ASE specifically in glutamatergic neurons (Extended Data Figs. 5b, 7l, and 8b), thereby complementing a previous bulk tissue study that characterized it as a mQTL.

Our approach to generating a high-confidence SV catalog and performing cell type-specific SV-eQTL and ASE analyses addresses a major gap in genetic studies, which have historically overlooked SVs. Conventional GWAS and resultant variant-to-function pipelines often focus predominantly on single-nucleotide variants (SNVs)^48^, despite the knowledge that SVs are a major source of genetic diversity and are disproportionately enriched for functional effects on gene regulation^27^. Short-read sequencing pipelines conventionally used in large studies often miss more than half of SVs^8,9^, particularly insertions and SVs in complex or repetitive regions. One of the current challenges in connecting variants to function is to elucidate what regulatory elements and cell types are affected. By combining epigenomic and genomic annotations with SVs, we can infer the former and by adding snRNA-seq data, we can understand the latter. In the future, this approach could be applied to other types of single-cell data sets as well as spatial transcriptomics.

Although our study brings new insight into the effects of SVs on gene expression, there are limitations inherent in applying these nascent technologies at the population scale. First, despite the advantages of long-read sequencing, structural variant genotyping accuracy remains lower than that of SNVs^8^. Not only is one SV caller insufficient for long-read sequence data, but genotyping tools applied subsequently also have limitations. Both Kanpig and VaPoR have limitations, which were shown by our benchmarking with synthetic data, in terms of SV type, length, and location within genomic regions with repeats (Supplementary Note 1). As such, larger and more complex SVs, such as the ∼900 kb inversion in the 17q21.31 region^49^, are less likely to be included in our high-confidence SV catalog. Second, the current cohort size of 100 individuals limits our statistical power to robustly assess the impact of rare structural variants and those with smaller effect sizes. While this sample size was likely sufficient to saturate the discovery of common SVs (AF ≥ 5%)^9^, assessing lower frequency variants requires more samples. Consequently, our current cohort size is not designed to robustly assess rare structural variants and their functional contributions to disease. Third, the power to detect SV effects on gene expression is lower in cell types present in our snRNA-seq dataset at lower frequencies, such as dopaminergic neurons in the midbrain (Supplementary Fig. 2d). Fourth, the sequencing depth (∼17x) and type of data in our study are suitable for a population-level approach, but not sufficient for accurate genome assembly. As has been described^50,51^, adding more HiFi reads, short reads, ultralong reads, and/or Hi-C data would be required, though at significant cost. Fifth, we used WASP^52^ to mitigate reference bias in snRNA-seq data, but it is based on the SNPs, so that we cannot rule out that some reference biases from SVs might be present in the ASE results. Despite these limitations, the integration of HiFi SV detection with cell type-specific functional genomics provides a valuable framework for identifying the effects of SVs on gene expression and prioritizing novel regulatory elements for functional follow-up.

## Methods

### Cohort

Our study included brain tissue from study participants the Arizona Study of Aging and Neurodegenerative Disorders (AZSAND) and the Brain and Body Donation Program (BBDP; www.brainandbodydonationprogram.org)^53^. Cases were selected to include the spectrum of PD. This spans from healthy controls to individuals with incipient, clinically asymptomatic Lewy body neuropathology, to those with clinically manifest PD. We included age- and sex-similar individuals in each of these three diagnostic groups: healthy controls (N = 34), ILBD (N = 31), and PD (N = 35) – see Lin et al. (manuscript submitted) for more details. ILBD cases were defined as individuals who did not meet clinical diagnostic criteria for PD or other neurodegenerative diseases but were found to have Lewy body inclusions at autopsy. ILBD can be considered a preclinical (prodromal) stage of PD. Cases designated as PD had both a clinical and a neuropathological diagnosis of Parkinson’s disease.

### Genomic DNA isolation

Cerebellum tissue samples were retrieved from -80°C storage and maintained on dry ice. Following manufacturer’s instructions using either the DNeasy Blood and Tissue Kit, (Qiagen, 69504) or the Nanobind PanDNA Kit (Pacific Biosciences, 103-260-000), genomic DNA was extracted (Supplementary Table 2).

For DNA extraction using the DNeasy Blood and Tissue Kit: Briefly, a 10-15 mg piece of tissue was cut and placed into a 1.5 ml tube with 180 μl Buffer ATL (Qiagen DNeasy Blood and Tissue Kit). DNA extraction proceeded following the Qiagen published protocol in the DNeasy Blood and Tissue Handbook: “Purification of Total DNA from Animal Tissues (Spin-Column Protocol),” starting from step 2 – Proteinase K digestion.

For DNA extraction using the PanDNA Kit: Briefly, a 5-10 mg piece of tissue was cut off using a disposable razor into a petri dish over wet ice. Then, 750μl cold Buffer CT (Pacific Biosciences PanDNA Kit) was added to the dish; tissue was disrupted via pipetting up and down 20 times, and entire volume was transferred into a 2 ml Protein LoBind tube (Eppendorf 0030108132). DNA extraction proceeded per the PacBio published protocol “Extracting DNA from Animal Tissue Using the Nanobind PanDNA Kit” (rev.03 MAR2024), starting from step 7 – first centrifugation step.

### Genomic DNA QC

Incoming genomic DNA (gDNA) samples were subjected to a quality check prior to library preparation for PacBio Circular Consensus Sequencing (CCS). This process involved a volume check, quantification using a Lunatic UV/Vis Spectrometer (Unchained Labs, 700-2000), and for a subset of samples, fragment analysis using a Femto Pulse gDNA assay (Supplementary Table 2.).

### Volume check and quantification

Sample volume was checked using a P200 pipette with filtered, wide bore tips (Fisher Scientific, 21-236-1A). For quantification, 3 μL aliquots of the samples were transferred from stock tubes (Nalgene B3732) to Lunatic plates (Unchained Labs, 701-2019) on a Bravo liquid handling platform (Agilent Technologies, G5509A) using Bravo Lab disposable 70 μL tips (Agilent Technologies, 19133-102). The Lunatic instrument was used to perform an A260 dsDNA (turbidity) assay to determine concentration. Total DNA mass was calculated using the concentration (ng/μL) and volume in the source tube. A minimum of 4 μg of DNA was required per attempt at SMRTbell Library Construction.

### Fragment analysis

Following the volume check and quantification, a subset of samples proceeded to sizing using the Femto Pulse gDNA assay (Supplementary Table 2). For those samples, fragment analysis was performed using the Genomic DNA165 kb Kit (Agilent Technologies, FP-1002-0275) on the Femto Pulse instrument (Agilent Technologies, M5330AA) according to manufacturer’s directions. Results were analyzed using ProSize software (v3.0; Agilent Technologies). Smear analysis was performed to calculate the percentage of DNA greater than 40 kb or greater than 10 kb in size. The specification for long-read sequencing required that more than 50% of DNA must be greater than 40 kb. If samples did not meet this criterion, a modified workflow was followed. Samples that had peaks above 23 kb and 15% of fragments ≥ 40 kb moved forward into shearing. Any sample outside the modified specifications skipped shearing and were manually transferred to a manually barcoded Abgene 96 Well 0.8mL Polypropylene DeepWell plate (Thermo Fisher Scientific, AB0859) targeting a mass of 4 μg in a volume of 110 μL.

### DNA shearing

A subset of purified genomic DNA (gDNA) samples (including the subset of samples with peaks above 23kb and 15% of fragments ≥ 40 kb and samples that did not undergo initial fragment analysis QC) underwent mechanical fragmentation to target a size distribution of 10 kb to 23.5 kb (Supplementary Table 2). A gDNA mass of 4 μg in a volume of 110 μL was transferred into barcoded 0.5 mL shearing tubes using a STARlet liquid handler (Hamilton Company, 173000). The DNA was fragmented on a Megaruptor 3 instrument (Diagenode, B06010003) using the Megaruptor 3 Shearing Kit (Diagenode, E07010003). A hydropore and syringe assembly was attached to each 0.5 mL shearing tube before being placed on the Megaruptor 3. Shearing was performed using the following parameters: Concentration: 49 ng/μL, Volume: 100 μL, and Speed: 31. After shearing, the DNA fragments were transferred from the shearing tubes to a barcoded Abgene™ 96 Well 0.8mL Polypropylene DeepWell™ plate using the Hamilton liquid handler. Post-shear, sample fragment distribution was evaluated using Femto Pulse as prescribed by the gDNA QC assay. Results were analyzed using ProSize software.

## SMRTbell library construction

SMRTbell Library Construction utilized the PacBio SMRTbell Prep Kit 3.0 (Pacific Biosciences, 102-182-700) and was fully automated on the STARlet liquid handler equipped with Hamilton filtered tips 50 μL (Hamilton Company, 235948), Hamilton filtered tips 300 μL (Hamilton Company, 235903), Hamilton filtered tips 1000 μL (Hamilton Company, 235905), a Magnum FLX with Solid-Core Technology magnet (Alpaqua, A000400), 37°C heat block, 67°C heat block, shaker, and tube chiller. Master mixes were made and added to the tube chiller. The library construction (LC) workflow consisted of a post-shearing cleanup, DNA repair, A-tailing, adapter ligation, and nuclease treatment, with cleanup steps performed after end repair/ligation and after nuclease treatment as detailed below. For post-shearing cleanup, a 1.0X volume over volume (v/v) of SMRTbell cleanup beads (Pacific Biosciences, 102-158-300) was added to the MIDI plate with sheared DNA. After a 30 second shake on the shaker block and a 10 minute incubation at ambient temperature, the MIDI plate was transferred to an Alpaqua Magnet and the supernatant was removed. 200 μL of 80% Ethanol was added to the plate, incubated for 30 seconds, and removed. This ethanol wash step was then repeated. The MIDI plate was then transferred off the Alpaqua magnet and 47 μL of 1X Low TE buffer (10 mM Tris-HCl, 0.1 mM EDTA, pH 8) was added. After a 30 second shake on the shaker block and a 5 minute incubation at ambient temperature, the MIDI plate was transferred back to the Alpaqua magnet. The eluate was then transferred to a Clear 96-well Plate (Eppendorf, 477-44-116). Following clean up, for DNA Repair and A-Tailing, 14 μL of DNA repair master mix (consisting of Repair buffer, End repair mix, DNA repair mix in a 4:2:1 ratio) was added to each DNA sample. After thorough tip mixing the 96-well plate was transferred to the appropriate heat blocks for DNA Repair and A-tailing thermal cycling (Step 1: 30 minutes at 37°C > Step 2: 5 minutes at 65°C). The sample then went through adapter ligation. For adapter ligation, 4 μL of SMRTbell adapters from SMRTbell adapter index plate 96A (Pacific Biosciences, 102-009-200) was added to the sample, followed by 31 μL of ligation master mix (consisting of ligation mix and ligation enhancer in a 30:1 ratio). After a 30 second shake on the shaker block the 96-well plate was transferred to the appropriate heat blocks for adapter ligation thermal cycling (30 minutes at 20°C). Following adapter ligation, a subsequent 1.0x v/v bead cleanup was performed using a 40 μL final elution volume. After clean up, the samples went through a nuclease treatment. For nuclease treatment, 10 μL of nuclease master mix (consisting of nuclease buffer and nuclease mix in a 1:1 ratio) was added to each DNA sample. After a 30 second shake on the shaker block, the 96-well plate was transferred to the appropriate heat blocks for nuclease treatment thermal cycling (15 minutes at 37°C). Following nuclease treatment, a subsequent 1.0x v/v bead cleanup was performed using a 25 μL final elution volume of 10 mM Tris-HCl pH 8.0 buffer. The final step of the automated LC protocol transferred the clean SMRTbell libraries to a matrix rack 0.64 ml tubes (Thermo Fisher Scientific, 3744PUR-BR).

### Post library construction QC and size selection

SMRTbell libraries were quantified using the Lunatic UV/Vis Spectrometer. Any sample that was not sheared skipped the size selection process to maximize the final loading concentration on the sequencer. Post-library construction quantification was reviewed for each of the sheared samples. The PippinHT’s historic size selection yield was calculated to be 42%. If the concentration of the sample was unlikely to yield enough material using this percentage, the PippinHT (Sage Sciences, HTP0001) protocol was skipped (Supplementary Table 2). Any sample with a high enough post-library construction yield was processed on the PippinHT to remove fragments less than 10kb. The PippinHT underwent an optical calibration by covering the LED detectors inside the machine with the manufacturer provided calibration fixtures. Once the calibration was complete, continuity testing was performed. The 0.75% Agarose PippinHT high-pass 6-10kb and 15-20kb cassettes (Sage Sciences, HPE7510) were brought to room temperature, prepped with electrophoresis buffer, and underwent continuity testing. This testing ensured that the cassette and current were adequate to run the PippinHT. After selecting “test,” the continuity test was run. After receiving a “pass” result, the cassette was loaded with the samples using a Bravo liquid handler (G5509A) and placed on the PippinHT. After loading the cassette onto the PippinHT, the “6-10kb High Pass Marker 75E” cassette definition was selected. The “BPstart” value was listed as 10,000bp, and the “BPend” value was listed as 50,000bp using the “Range” option. After the marker lane was identified, the size selection process was started. Size selection ran at room temperature for approximately 1 hour and 20 minutes at 75V. Following the run, cassettes were allowed to sit for 45 minutes at room temperature in an effort to increase yield before the samples were unloaded by the Bravo liquid handler (G5509A) using 0.1% Tween20 in TE pH 8.0 (0.1% Tween 20, 10 mM Tris, 0.1 mM EDTA) as a wash. The recovered samples underwent a final 1.0x v/v SMRTbell cleanup (95 μL of SMRTBell cleanup beads) on the STARlet liquid handler as described in “SMRTBell Library Construction” to purify size selected libraries. The cleaned post-PippinHT volume (23 μL) was transferred to a new matrix rack containing 0.5 mL tubes (Thermo Fisher Scientific, 3744BLU-BR).

### Final library quantification, normalization, and pooling

The final cleaned libraries were quantified again using the Lunatic prepared with the Bravo liquid handler (G5509A), as was performed post LC. Samples were then normalized using EB (10 mM Tris HCl pH 8.0) to target a final sequencing loading concentration of 215-250pM before entering the Sequencing Prep workflow (Supplementary Table 2 for details). Libraries were transferred to new matrix racks (Thermo Scientific, 3744BLU-BR) for downstream sequencing preparation. In the event that a library did not yield enough material after library construction or size selection, the entire recovered library was used.

### PacBio automated annealing, binding, and cleanup (ABC)

SMRTbell libraries were transferred from matrix racks to 96-well plates (Eppendorf, 477-44-116) and underwent annealing, binding, and cleanup (ABC) using automated Hamilton protocols on the STARlet. Initially, SMRTbell cleanup beads (Pacific Biosciences, 102-158-300) and ABC buffers were brought to room temperature, while sequencing primer, internal control, and polymerase (from the Revio Polymerase Kit, Pacific Biosciences, 102-739-100) were thawed on ice. A 25 μL aliquot of master mix containing sequencing primer and buffer (1:1) was added to each sample to hybridize the sequencing primer to the SMRTBell template and allowed to incubate for 15 minutes at ambient temperature. Upon completion of primer hybridization, 1 μL of polymerase and 49 μL polymerase buffer were added to each sample, and allowed to incubate for 15 minutes. Subsequent to the binding reaction, 120 μL of SMRTBeads were added to the sample volume for a 1.2X Ampure PB SPRI Clean Up. Bead binding occurred over a 10 minute incubation at ambient temperature before moving to the Alpaqua magnet. Once beads had precipitated out of solution, supernatant was removed and beads were eluted in 100 μL of Revio loading buffer (from the Revio Polymerase Kit, Pacific Biosciences, 102-739-100) and allowed to bind for 5 minutes at ambient temperature. Libraries were once again moved to the Alpaqua magnet and once beads had precipitated out, final bound complexes were transferred to a clean 96-well PCR plate (Eppendorf, 477-44-116). This cleanup step was performed without an ethanol wash. Following purification, the Revio Sequencing Control (Pacific Biosciences, 103-508-800) was manually diluted following PacBio’s guidelines and spiked into the final clean complexes. Samples were stored in the dark at 4°C for up to 24 hours prior to sequencer loading. All ABC steps were conducted using automated protocols, with specific volumes determined by the SMRT Link software for each sample (Supplementary Table 2 for version numbers).

### Sequencing plate preparation and instrument loading

Upon completion of the ABC process, samples were prepared for loading onto the PacBio Revio sequencing instruments. Revio Sequencing Plates were thawed from -20°C. Each kit included a sealed plate and a unique QR code card, which were maintained as matched sets. Plates and QR codes were labeled with corresponding identifiers for tracking. Automated transfer of post-ABC samples into the Revio sequencing plates was performed on the Bravo liquid handling platform (Agilent Technologies, G5563A), using 180 µL Bravo tips (Agilent Technologies, 08585-002). The pierced wells of the reagent plate were manually sealed with foil seals (Pacific Biosciences, 100-667-400) provided within the kit.

Run design creation and upload to SMRT Link were automatically generated by a Broad Institute laboratory information management system. The Revio sequencer was loaded as described with the manufacturer protocol (Pacific Biosciences, PN 102-962-600 REV03). Sequencing was performed using 25M Revio SMRT Cells (Pacific Biosciences 102-202-200). Each SMRT cell was run with a 2-hour pre-extension time and a 24-hour movie time to generate raw sequencing data. The processing of these samples spanned two versions of SMRT Link software (Supplementary Table 2 for details).

### Long-read data processing

We used workflows included in the Broad Institute long-read (LR) pipelines^54^ (v4.0.58) to process PacBio HiFi WGS data. These pipelines are written in WDL^55^ (v1.0) and executed on the Terra cloud platform^56^. By applying PBFlowcell workflow, raw subreads of each sample were first processed into consensus HiFi reads using CCS^57^ (v6.2.0). The workflow then aligned these reads to the GRCh38 human reference genome using pbmm2^58^ (v1.4.0), and the resulting alignments were merged and sorted with SAMtools^59^ (v1.13) to produce a single, aligned BAM file per individual.

### Individual-specific SV detection

To ensure high sensitivity and robustness in variant discovery, we employed an ensemble SV calling strategy using three distinct algorithms on each individual’s aligned BAM file. Brief information about each variant caller used along with invoked parameters is listed below:

- **Sniffles2**^23^ (v2.0.6) is an alignment-based method that extracts SV signatures from read alignment and calls SVs based on engineered set of rules, such as split reads and read depth. This caller was run as part of the PBCCSWholeGenome workflow in Broad Institute’s LR pipeline with the default parameters on Terra cloud platform.
- **PBSV**^22^ (v2.9.0) is PacBio’s official suite of tools used to call and analyze SVs in diploid genomes from PacBio single molecule real-time sequencing (SMRT) reads. This caller is another alignment-based method that first discovers SV signatures from read alignment, creating SVSIG files. During this “discover” phase, tandem repeats BED file (for GRCh38 no alt^60^ downloaded on October 31, 2023) was provided to increase sensitivity and recall as recommended by the authors. This caller was also run as part of the PBCCSWholeGenome workflow in Broad Institute’s LR pipeline with the default parameters on Terra cloud platform.
- **CUE**^21^ (v2.0.0) is a deep learning-based caller that calls deletions, duplications, inversions and inverted duplications. Cue2 is an extension of the image-based deep learning framework Cue, optimized for long reads. It includes new image channels designed specifically for long reads and a new module to recover SV signals from pervasive alignment artifacts. The pre-trained Cue2 HiFi model was downloaded from [https://storage.googleapis.com/cue-models/hifi/cue2.hifi.v1.pt]. Cue2 was run on the Broad Institute’s high-performance computing cluster with “n_cpus” parameter set to 16. All other parameters were left as default.

This multi-caller approach generated three separate VCF files (one from each caller) per individual, capturing a comprehensive initial set of SVs.

### Cohort-level SV processing and filtering

The initial individual-specific SV callsets generated by the ensemble strategy were processed through a multi-stage pipeline to create a single cohort-level VCF for the entire PD5D cohort. First, low-quality variants (VCF FILTER column is not “PASS”) and those smaller than 40 bp were removed from all VCFs using the InitialFilterVCF^61^ Python script (Python (v3.9.23)) to facilitate SV matching. To prevent the erroneous merging of different SV types, the calls were split by SV type (insertion, deletion, duplication, and inversion) using the BCFtools^59^ (v1.21) view command with -i parameter. The 100 individual-specific VCFs of matching SV types from each caller were then combined into a single multi-individual VCF using the SURVIVOR^24^ (v1.0.7) merge command with the following parameters: maximum allowed distance of 1 kb, minimum number of caller support of 1, require SV strand match, and minimum SV size of 50 bp. Subsequently, the resulting three VCFs (one from each caller) for each SV type were merged using SURVIVOR (v1.0.7) with the same parameters. Finally, the resulting four VCFs (one in each SV type) were combined using the BCFtools (v1.21) concat command with -a parameter, yielding a single VCF for the PD5D cohort.

Next, we excluded SVs based on the genomic location, removing SVs in centromeres, reference genome assembly gaps, and known segmental duplications (downloaded from UCSC table browser^60^ on August 19, 2024 (centromeres), July 8, 2025 (assembly gaps), and November 4, 2025 (segmental duplications)). These genomic intervals were combined using BEDTools^62^ (v2.29) and then overlapping SVs were filtered using BCFtools (v1.21) filter method specifying the target file. We also removed SVs on the sex chromosomes (X and Y) due to the unequal representation in males and females.

We then performed read-level quality control to exclude low-confidence SVs (Extended Data Fig. 1g). For each SV in the VCF, we applied the AddSVQC^63^ Python script (Python (v3.9.23)) to load the VCF file using cyvcf2^64^ (v0.31.4) to first identify carrier samples (both HET and HOM ALT) in each diagnostic groups (HC, ILBD and PD). Treating HC as a control group and ILBD and PD as a case group, the frequency of the SV in each group (CASE_RATE and CONTROL_RATE) along with carrier sample IDs (HC_CARRIERS, ILBD_CARRIERS and PD_CARRIERS) were annotated. The script then used pysam^65^ (v0.23.3) to extract reads found in the genomic interval of the SV from carrier sample BAM files and annotated SVs with the total number of reads across all carriers (TOTAL_READS), average read depth per carrier (AVG_READS_PER_SAMPLE), and average mapping quality of those reads (AVG_MAP_QUALITY). Lastly, combining information from the SNP genotype array (PD5D MEGA chip genotype dataset available in the CRN cloud “scherzer-pmdbs-genetics”; DOI 10.5281/zenodo.17242295), the script calculated r^2^ ^66^ to quantify linkage disequilibrium (LD) between the SV genotype and neighboring SNP genotypes. This calculation was restricted to SNPs within a flanking distance of 10 kb of the SV breakpoints. We counted the total number of nearby SNPs (TOTAL_SNPS_NEARBY) and those that achieved high-LD (r^2^ ≥ 0.7) (LD_SNPS_COUNT). The ratio of high-LD SNPs to total nearby SNPS was also reported (LD_SNP_RATE). Using these read-level quality metrics, we excluded low-confidence SVs with < 5 TOTAL_READS, average number of reads across carrier samples < 5, average mapping quality of supporting reads < 20 (Phred score), or alternate allele count < 1. Note, SV genotypes used to calculate these metrics were assigned following our strategy explained below.

### Comparison of SV callsets from the three callers

We used Truvari^35^ (v5.3.0) to compare the initial combined callsets from different callers. Truvari was selected because its systematic approach leverages multiple similarity metrics, including sequence similarity, which is essential for accurately comparing different alleles sharing the same or similar breakpoint coordinates. First, Truvari collapse (with –dup-to-ins) was run on each callset to allow for matches between insertions and duplications and merge potentially redundant calls. Then, using Truvari’s bench module (with --reference, --sizemax=100M, --sizefilt=30, --sizemin=50, --refdist=500, --dup-to-ins), we compared callsets between Cue2 and Sniffles2, and between Cue2 and PBSV (predictions from Cue2 were used as the base callset in both comparisons). Next, the remaining unmatched variants from Sniffles2 and PBSV were similarly compared, using the Sniffles2 callset as the base callset. The comparison results were then converted to a Python loadable file using Truvari’s vcf2df module (with --info, and --bench-dir). We applied the PrepUpsetData^67^ script (Python (v3.9.23), joblib^68^ (v1.5.3), pandas (v.2.3.1), NumPy^69^ (v2.0.2)) to collect a list of matching variant IDs and count unique SV type combinations. The resulting counts were used to create an upset plot using the MakeUpset^70^ script (R (v4.5.1) and R package ComplexUpset^71^ (v1.3.3)).

### SV annotation

To assign functional, regulatory, and clinical relevance to the final set of SVs, we used AnnotSV^20^ (v3.5.3). We ran AnnotSV using the following parameters: SVinputFile, outputFile, REreport=1, missingGTinSamplesid=0, genomeBuild=GRCh38 and vcf=1. The relevant annotations provided in this analysis are listed:

- Gene/Region-based annotations: RefSeq genes^72^ (accessed July 17, 2025). the Gene Curation Coalition (GenCC)^73^ (version date April 22, 2025). Online Mendelian Inheritance in Man (OMIM) morbid genes^74^ (version date June 5, 2025). Haploinsufficiency^75^. Cytoband^76^ (downloaded July 17, 2025). GC content +/- 100 bp from each SV breakpoint^77^ (downloaded July 17, 2025).
- Regulatory elements: Promoters (500 bp upstream from the transcription start sites of the RefSeq database (accessed July 17, 2025)). Enhancers reported in EnhancerAtlas 2.0^78^. Promoters/enhancers reported in GeneHancer^34^. miRNAs reported in miRTargetLink 2.0^79^. Predicted enhancer-gene regulation by activity-by-contact model^80^. Functional measurements from massively parallel reporter assays on 20 disease-associated gene promoters and enhancers^81^. Among these data sources, GeneHancer data were specifically curated to select brain-active regulatory elements using the linked tissue expression data (see GitHub). All brain-related tissues were included (tissues with a case-insensitive pattern “brain”, and other brain related tissues: meninx, olfactory region, astrocyte, cranial nerve, dorsal root ganglion, neural progenitor cell, neural stem cell, neural stem progenitor cell, neural tube, neuron, neuronal stem cell, and spinal cord) and GeneHancer activity score (“combined_score”) > 10 was considered active.

AnnotSV result also included SV ranking score based on overlapping known benign/pathogenic regions reported in gnomAD^82^, ClinVar^83^, ClinGen^84^, the Database of Genomic Variants^85^, DECIPHER^86^, 1000 Genomes Project (1KGP) Phase 3^87^, NCBI Curated Common Structural Variants (nstd186)^88^, a disease trait-mapping study^46^, Children’s Mercy Research Institute (CMRI) database^89^, Human Pangenome Reference Consortium (HPRC)^51^, CoLoRSdb^90^. AnnotSV predicted pathogenicity is reported following the ACMG (American College of Medical Genetics) recommended^91^ annotations. Details are in Supplementary Table 6.

We additionally cross-referenced SV positions in the genome with other information available in the UCSC database for GRCh38^92^, including transcription factor binding sites (encRegTfbsClustered), DNase hypersensitivity sites (wgEncodeRegDnaseClustered), CpG islands (cpgIslandExt), evolutionarily conserved sites (phastCons100way), and H3K4me1 (wgEncodeBroadHistoneH1hescH3k4me1StdSig) and H3K27ac (wgEncodeBroadHistoneH1hescH3k27acStdSig) sites in the H1 cell line. The proportion of SVs overlapping these sites along with statistical significance was assessed in R (v4.2.2) via permutation testing (permTest function) implemented in R package regioneR^93^ (v1.30.0) with 1,000 iterations. To account for genomic biases, we defined an "allowed universe" by masking gaps, centromeres, and segmental duplications. We employed a custom resampling strategy that randomized SV locations while strictly preserving original segment widths and weighting by available block sizes. For each feature, we calculated fold enrichment and P-values by comparing observed overlaps against the permuted null distribution.

### Sequence analysis using Dfam

To characterize the repetitive content and origin of the identified SVs, specifically for the examples we describe in the text, we performed a homology-based search against the Dfam database^94^, using an online Dfam sequence search tool (v3.9) (https://www.dfam.org/search/sequence). Organism was selected to “Homo sapiens” and “Dfam curated threshold” was selected as cut-off. Inserted/deleted sequences of the eight SVs (chr8_22625880_INS_330, chr12_123521586_DEL_-108, chr12_123581929_INS_974, chr10_15531400_INS_60, chr14_55489789_DEL_-65, chr4_76150170_DEL_-109, chr17_46237501_DEL_-724, and chr17_46252361_INS_828) were provided in a FASTA format.

### Benchmarking SV concordance using public databases

The comparison of the identified SVs leveraged nine large-scale catalogs of genomic variation including two callsets from HPRC (with distinct variant calling methods: PBSV^22^ (also used in this study) and an assembly-based method^95^), CoLoRS, CMRI, deCODE^7^, gnomAD, 1KGP Phase 3, disease trait-mapping, and ClinVar. Detailed information about each database, including sequencing technology, variant calling strategy, cohort size, available SV types, and dataset link are available in Supplementary Table 6.

To query each database and assess the precision and recall of our SVs, we used Truvari^35^ (v5.3.0). We used Truvari’s bench module to test two distinct parameter settings against each public catalog: strict and lenient. Strict setting only allows near-perfect matching by setting the maximum allowed reference distance (--refdist) of 1 bp, and requiring sequence similarity (--pctseq), size similarity (--pctsize), and reciprocal overlap (--pctovl) >99%. By contrast, the lenient setting uses the default settings of the tool for a more relaxed comparison (a 500 bp breakpoint window and >70% similarity across size, overlap and sequence.

### Synthetic SV data

Synthetic datasets used to evaluate genotyper performance were obtained from the Varium repository (https://github.com/PopicLab/varium). These datasets were designed to systematically stress-test SV recall and precision as a function of SV type, size, and genome context. We downloaded 60 datasets spanning four SV types (DEL, INS, DUP, and INV), five SV size regimes (50-150bp, 150-500bp, 500-1kb, 1k-10kb, and 10k-20kb), and three genomic contexts (MAPPABLE, NONUNIQUE, and SEGDUP). As defined in the repository README, MAPPABLE regions are defined as a complement of all the RepeatMasker tracks^96^, while the NONUNIQUE and SEGDUP regions correspond to the Genome in a Bottle (GIAB) mappability stratifications provided in the GRCh38_nonunique_l250_m0_e0.bed.gz^97^ and GRCh38_segdups.bed.gz^98^ BED files, respectively. Each dataset is derived from a synthetic genome with 1000 simulated SVs of a particular type and size placed in a specific genome context using insilicoSV^99^. For each dataset, we downloaded the 30x BAM file with synthetic HiFi reads and the truth set VCF file (with ground truth SVs).

### Genotyper performance benchmark

We benchmarked the performance of Kanpig^37^ (v1.1.0) and VaPoR^38^ (no version number) on the 60 synthetic HiFi datasets with SVs of different type and size in three sequence repeat contexts. Kanpig was run on Broad Institute’s HPC to predict genotypes of all simulated SVs for each SV type with the following parameters: --hapsim 0.97, --chunksize 500, --maxpaths 1000, and --gpenalty 0.04. On the same datasets, for inversions only, we additionally benchmarked VaPoR. VaPoR was run on the Terra cloud platform using the WDL included in the GATK-SV pipeline^100^ (v1.1). Dockers used for different tasks are listed: us.gcr.io/broad-dsde-methods/gatk-sv/sv-base-mini:2024-10-25-v0.29-beta-5ea22a52 to subset VCF to a sample and to concatenate VaPoR results, us.gcr.io/broad-dsde-methods/gatk-sv/sv-pipeline:2025-10-02-v1.1-483973d6 to convert VCF to BED and to extract specific contig from BED, and us.gcr.io/broad-dsde-methods/eph/vapor:header-hash-2fc8f12 to run VaPoR. These results were downloaded from the Terra cloud platform for the downstream comparison to the ground truth using Truvari (v5.3.0) bench module with the default parameters and --reference. The genotype concordance results across SV types and sequence repeat contexts were collected and plotted using R (v.4.2.2) and R package ggplot2^101^ (v3.4.4).

### Genotyping SVs

Insertions, deletions, and duplications within the above defined confident size ranges were selected and genotyped using Kanpig. Kanpig was used rather than the native genotyping modules of individual callers to ensure consistency and because it was shown to achieve higher genotyping accuracy than Sniffles2 (and several tools not included in this study), especially when there are multiple neighboring SVs^37^. Kanpig was run on the Terra cloud platform (docker: quay.io/zhengxc93/terra-kanpig) with the identical parameters as used for benchmarking. Genotypes for inversions were left unchanged, as the genotyper performance evaluation revealed no suitable tool for accurate annotation of inversions genotypes, and were all included in subsequent analyses. For those inversions with conflicting genotypes calls, PBSV genotypes were used when available, and Sniffles2 otherwise, given the lack of public Cue2 genotyping-focused benchmarks..

### SV selection for functional analysis

We used a targeted strategy by focusing the functional analysis on SVs located within 100 kb of the previously identified 124 PD GWAS eGenes (Lin et al., manuscript submitted). To maintain statistical power in the small cohort (N=100), we excluded rare SVs with an alternate allele count < 2, those with valid genotype calls in < 5 samples (./. is not valid), monomorphic SVs with allele frequency ≥ .99, and those detected only in healthy control individuals. This filtering was performed using BCFtools (v1.21) filter module with -i parameter (bcftools filter -i ’F_MISSING <= 0.95 && AF < .99 && (AC_PD > 1 || AC_ILBD > 1)’). Sample group-level allele counts were added to SVs using BCFtools (v1.21) fill-tags module with -S parameter to provide diagnostic group information.

### Hybrid selection of snRNA-seq midbrain libraries

Hybrid selection for PD midbrain snRNA-seq libraries was performed based on our previously described method^12^, with several modifications. To obtain sufficient input material, each individual library was amplified using P5 (5’-AATGATACGGCGACCACCGA-3’) and P7 (5’-CAAGCAGAAGACGGCATACGA-3’) primers and Amp Mix (#2000047, included in the 10x Genomics BCR Kit). Each 50 µL PCR reaction contained 25 ng of library, 25 µL Amp Mix, 0.5 µL of 10 µM P5, 0.5 µL of 10 µM P7, and nuclease-free water to volume. Libraries were amplified for 6 PCR cycles using the following program: 45 seconds at 98°C; 6 cycles of 20 seconds at 98°C, 30 seconds at 60°C, and 20 seconds at 72°C; followed by 1 minute at 72°C and hold at 4°C. PCR products were purified using 0.8× AMPure XP SPRI beads (Beckman Coulter, A63881) with freshly prepared 80% ethanol.

The 96 midbrain libraries were then combined into 12 pools with balanced cell numbers and index representation. Hybrid selection was performed using the SureSelect Max Overnight Hyb Kit (Agilent, G9690A) according to the manufacturer’s instructions with the following modifications: (1) 750 ng of pooled library was used as input; (2) 1 µL each of universal blocking oligos TS-p5 and TS-p7 (IDT, 1016184 and 1016188) was added during the denaturation step; (3) hybridization was performed at 65°C overnight (∼20 hours); and (4) post-capture PCR amplification was performed for 14 cycles.

For sequencing, post-capture products were combined into two “super pools,” each containing samples with non-overlapping indices. Final pooled libraries were sequenced on a NovaSeq X Plus 25B (Illumina, 20084804) using a 300-cycle kit (Illumina, 20104706) with the following configuration: 28 bases for Read 1, 150 bases for Read 2, 10 bases for Index 1, and 10 bases for Index 2.

### snRNA-seq data for function analysis

We used the MTG and midbrain snRNA-seq dataset available in the ASAP Collaborative Research Network (CRN) Cloud in the “ASAP Parkinson Cell Atlas in 5D (PD5D)” dataset (DOI: 10.5281/zenodo.16751625, see the dataset “team-scherzer-pmdbs-snrnaseq-mtg” and the link to midbrain snrna (DOI in progress)). Sample-level pseudobulk expression data were prepared following the procedures and quality control detailed in Lin et al. (manuscript submitted), with details of the pseudobulking explained below. We used the 94 out of 100 samples found in both the VCF and snRNA-seq data in each brain region. The following samples were not included due to issues with snRNA-seq data such as low quality, sample tracking, or being a genotype outlier: BN1610, BN1750, BN1864, BN1867, BN1975, and BN9950 in the MTG and BN0009, BN0602, BN0635, BN0704, BN1022, and BN1610 in the midbrain.

For the pseudobulking, after normalizing the count data (R (v4.2.2), Seurat (v4.3.0) NormaliseData function, params: normalization.method = "LogNormalize", scale.factor = 10000), the average expression of genes of all cells from a cell type were investigated and the effect of covariates including sequencing batch, sex, age, RIN, PMI and hidden relatedness among individuals were adjusted as follows. Batch effects were first adjusted using the ComBat^102^ function of surrogate variable analysis (SVA) package^103^ (v3.46.0) in R (v4.2.2). Adjusted gene expression values were then used as input for the sva function to identify hidden confounders. A linear model implemented in the fsva function of SVA package was used to adjust for the known covariates of age, sex, RNA integrity number, and post-mortem interval as well as hidden covariates captured in the first 30 SVA dimensions. Finally, the covariance adjusted pseudo-bulk count data was scaled using the scale function included in R base package.

### *cis*-SV-eQTL mapping

The curated VCF was first processed using the prep_SVs^104^ R script (R (v4.2.2)) to generate specific input files required for a MatrixEQTL (R package version 2.3) analysis. It converts raw genotype strings (e.g., “0|1”) into a numeric dosage matrix (0, 1, 2) and calculates SV end positions based on the SVLEN info field, generating two tab-separated output files: one for genotypes and the other for SV genomic positions. We then performed *cis-*eQTL analysis using MatrixEQTL^105^ (v2.3) across distinct brain cell types in the MTG (astrocytes, endothelial cells, fibroblast-like cells, GABAergic neurons, glutamatergic neurons, microglia, oligodendrocytes, oligodendrocyte precursor cells and pericytes) and in the midbrain (astrocytes, dopaminergic neurons, endothelial cells, fibroblasts, GABAergic neurons, microglia, monocytes, oligodendrocytes, oligodendrocyte precursor cells and pericytes). Benjamini–Hochberg false discovery rate (FDR) ≤ 0.05 was used as the significance cutoff. Bubble plots were generated with the ASE_eQTL_bubbleplot^106^ script using R package ggplot2^101^ (v3.4.4).

To compare the baseline eQTL results to one that additionally accounts for population stratification, we added six genotype principal components (PCs) as additional covariates and ran the above model again (cvrt parameter with six genotype PCs in rows). The genotype PCs were calculated using SNPRelate^107^ (v1.32.2) in R (v4.2.2). The SNP VCF file was first converted to genomic data structure (GDS) format (gdsfmt R package (v1.34.1)) using the snpgdsVCF2GDS function. After pruning SNPs with snpgdsLDpruning function (with parameters autosome.only=T, maf=0.05, missing.rate=0.05, method=’corr’, slide.max.bp=10e6, and ld.threshold=sqrt(0.1)), genotype PCs were computed with snpgdsPCA (with parameter algorithm=’randomized’).

To better understand SV-eQTLs that showed significance in only one cell type, we first established a binary expression matrix to identify cell types where eGenes were confidently expressed (> 50% of samples with mean UMI counts > 0.1). Then, a cell number cutoff (median number of cells across individuals >= 9) was used to identify cell types with limited power. Three cell types in the midbrain (dopaminergic neurons, fibroblast, and glutamatergic neurons) did not meet this cell number cutoff. For every significant SV-gene pair identified as significant in only a single cell type at the baseline level, we applied a four-way classification logic to categorize the basis of its specificity:

- Low expression: The effect is significant in only one cell type and the target gene is only confidently expressed in that single cell type across the entire region.
- Low cell number: The gene is expressed in multiple cell types, but the other cell types expressing the gene did not meet the minimum cell number cutoff.
- Low expression and cell number: The gene is not confidently expressed in other cell types nor did the gene meet the minimum cell number cutoff for those other cell types.
- Potential regulatory specificity: The effect is significant in only one cell type and the target gene is reliably expressed (meeting both UMI and cell count thresholds) in two or more cell types.

### Phasing SV data and ASE

To phase our SV data, we took the SV VCF we produced and a phased SNP level VCF on these same individuals that has been previously published^12^ (PD5D MEGA chip genotype dataset; DOI 10.5281/zenodo.17242295). We built a pipeline (PhaseLongRead, https://github.com/seanken/PhaseLongRead) with WDL to perform phasing for each sample in the VCF. The pipeline begins by preparing both VCFs with BCFtools^59^ (v1.11; using staphb/bcftools:1.11 as the Docker image), which involves sorting (bcftools sort) and indexing (bcftools index) the VCF, extracting the sample of interest (bcftools view with the -s flag), and filtering out non-heterozygous locations (bcftools view with the -g het flag). The BAM file for the long-read data is then split into three BAM files—reads coming from allele 1, allele 2, and those that are uncertain. To do this we created a jar file, written in Java (v11) (using the Docker image at amazoncorretto:11), which is a modified version of a tool we previously developed for long-read snRNA-seq ASE^12^ based on htsjdk^59^ (v2.21.1; see https://github.com/seanken/ASE_pipeline/tree/main/src/Java for the original tool). First a HashMap was created, mapping from each allele of each SNP (a class created for this implementation) to Boolean values (true representing if the allele is on allele 1, false if on allele 2). Each read was processed one by one, excluding those with supplemental mappings. The position in the genome and nucleotide for each position in the read were extracted, and the SNP VCF was used to determine if it matched a SNP from the SNP VCF and, if so, which allele it matched. If > 95% of SNPs in the read matched a given allele, the read was saved to the corresponding BAM, otherwise it was discarded. This resulted in the 3 BAMs described above. We then process the allele 1 and allele 2 BAMs separately (using the docker image amazoncorretto:11). We prepared these BAMs with SAMtools^59^ (v1.19) (using staphb/samtools:1.19 as the Docker image), by filtering them (with samtools view -b -- expr ‘[AL]>3 || [AM]>3’) and indexing them (with samtools index). We then ran Sniffles (v2.0.6) (using us.gcr.io/broad-dsp-lrma/lr-sniffles2:2.0.6 as the Docker Image) on each of these BAMs, using the --genotype-vcf flag to pass in the prepared SV VCF from above. This resulted in an SV VCF for allele 1 and one for allele 2, where allele 1 and allele 2 are based on the phased SNPs. Finally, these two VCFs were combined with Python (v3.14.3) (using python:3 as the docker image). First, both VCFs were loaded as DataFrames with pandas^108^ (v2.3.3). The genotype was extracted for each SV (based on the 0/0, 1/1, etc., values in the VCF). Those with 0/0 for allele 1 and 1/1 for allele 2 were assigned as 0/1 genotypes, those with 1/1 for allele 1 and 0/0 for allele 2 were assigned as 1/0, and all others were assigned as ./. since those are ones that we could not phase successfully. The results were then saved as a VCF with pandas (with no header), with one such VCF per individual. These VCFs were downloaded and added to an existing metadata file with one entry per cell, which contains cell type labels, disease status, and containing file paths to the outputs produced for performing SNP based ASE on these same samples^12^. We then used scAlleleExpression^12^ (v1.0) to load the pseudobulked ASE information for all cells/samples for the SV/gene pairs of interest with the GetSNPs command using these metadata. For each cell type, we then performed allelic imbalance analysis with the TestSNP_aod command.

## Data and code availability

PD-related WGS data will be available on Zenodo (DOI: 10.5281/zenodo.19124632). The associated genetics dataset (scherzer-pmdbs-genetics) is available on Zenodo (DOI: 10.5281/zenodo.17242295). Single-nucleus RNA-seq datasets for the MTG are available on Zenodo for both the original dataset (DOI: 10.5281/zenodo.16751625) and the hybrid-selected dataset (DOI: 10.5281/zenodo.16885839). The snRNA-seq datasets for the midbrain will be available on Zenodo for both the original (DOI: <u>10.5281/zenodo.18989318</u>) and the hybrid-selected dataset (DOI: 10.5281/zenodo.19124469). Initial long-read sequencing data processing was performed using the Broad Institute’s long-read pipelines (v4.0.58), available on GitHub (https://github.com/broadinstitute/long-read-pipelines). All custom code generated for this manuscript is deposited on Zenodo (DOI: 10.5281/zenodo.18676281). Additionally, the custom code used specifically for ASE analysis has been made available on Zenodo, including the PhaseLongRead pipeline v1.0 (https://zenodo.org/records/18509861) and the scAlleleExpression R package v1.0 (https://zenodo.org/records/10697254).

Supplementary Table 11 is provided as a key resource table.

## Acknowledgements

For the purpose of open access, the author has applied a CC BY public copyright license to all Author Accepted Manuscripts arising from this submission. We thank Felecia Cerrato for project management, Steve Huang for guidance on SV calling, Daniel El Kodsi for organizing genomic DNA transfers, Beatrice Weykopf for project management and assistance with open science compliance, Leslie Gaffney for assistance in drafting figures, and the Broad Institute’s Genomic Platform for DNA sequencing. We are especially grateful to Erin LaRoche, Stephanie Durgin, and Michael Dasilva for contributing text for the PacBio sequencing lab methods. We gratefully acknowledge the generous participants of the Arizona Study of Aging and Neurodegenerative Disorders (AZSAND) who donated their brains through the Brain and Body Donation Program. The samples provided by these participants were essential for the completion of this study.

## Funding

This research was funded by Aligning Science Across Parkinson’s [grant # ASAP-000301 and ASAP-024434] through the Michael J. Fox Foundation for Parkinson’s Research (MJFF) (J.Z.L., X.D., C.R.S), the Yale School of Medicine’s Stephen & Denise Adams Center for Parkinson’s Disease Research (C.R.S.), the Yale American Parkinson Disease Association Center for Parkinson Precision Medicine (C.R.S.), the Lineberger Research Fund (C.R.S.), NIH R01NS115144 (C.R.S), the U.S. Department of Defense (C.R.S), and NIH R01HG012467 (V.P.). The Brain and Body Donation Program has been supported by the National Institute of Neurological Disorders and Stroke (U24 NS072026 National Brain and Tissue Resource for Parkinson’s Disease and Related Disorders), the National Institute on Aging (P30 AG019610 and P30AG072980, Arizona Alzheimer’s Disease Center), the Arizona Department of Health Services (contract 05700, Arizona Alzheimer’s Research Center), the Arizona Biomedical Research Commission (contracts 4001, 0011, 05-901 and 1001 to the Arizona Parkinson’s Disease Consortium) and the Michael J. Fox Foundation for Parkinson’s Research.

## Author contributions

C.R.S. and J.Z.L. conceived the study. G.E.S selected the cases. M.P.C. collected the samples. T.G.B. obtained funding (for the cohort) and performed neuropathological characterization. Z. Liao and M.K isolated and provided genomic DNA. I.T., Z. Liao, and M.S. extracted and quality-controlled RNA, captured nuclei, generated snRNA-seq libraries, performed QC, and generated snRNA-seq data. J.P. processed, quality-controlled, and analyzed the snRNA-seq data. N.H. and J.Z. performed hybrid selection with snRNA-seq libraries. K.K. performed all the SV computational analysis, except as noted. Z. Liu and X.D. provided expert guidance on SV analysis. S.K.S. performed the allele-specific expression analysis, built the SV phasing pipeline, and advised on other analyses. V.P. advised on SV analysis. J.Z.L. directed the study and guided the analysis. All authors contributed to writing the manuscript, read, and approved the final version.

## Competing interests

All authors declare no competing interests.

## Extended Data Figures

**Extended Data Fig. 1 | Sequencing QC metrics**. Box plots showing the quality of HiFi reads and alignment. Samples are grouped based on clinical diagnosis (x-axis). Data points show values from each sample. The metric visualized is shown by the title in each subplot with the values on the y-axis.

**Extended Data Fig. 2 | Computational analysis pipeline. a**, Graphical summary of individual-specific SV calling pipeline. **b**, Graphical summary of SV processing pipeline. Colors highlight key steps: SV removal (red), SV combining (green), and SV annotation (yellow).

**Extended Data Fig. 3 | SV properties. a**, Bar chart showing the top 30 cytobands with the most SVs found. The number of SVs is shown on the x-axis and the cytobands are shown on the y-axis. Exact numbers are written on the right side of the bars. **b**, Upset plot showing the number of SVs overlapping with regulatory elements annotated by different sources. The bubble plot lists sources of regulatory elements and shows set memberships. Black circles indicate which sets are included in a particular intersection. Upper bar chart shows the size of each intersection. Set size bar chart (right) shows the total number of SVs in each individual set. The intersection bar plot is color-coded by SV type. Source acronym: EA=Enhancer Atlas, ABC=Activity-by-contact model, HI=Haploinsufficiency, GH=GeneHancer, mTL= miRTargetLink 2.0, RefSeq=NCBI reference sequence. **c**, Bar chart showing the observed and expected percentage of SVs overlapping with annotated genomic features. Features include regulatory elements from GeneHancer, brain-active regulatory elements from GeneHancer, CpG islands, DNase hypersensitive sites, H3K27ac marks from the H1 hESC line, H3K4me1 marks from the H1 hESC line, evolutionarily conserved sites from 100 vertebrate species, repeat regions identified by RepeatMasker, and transcription factor binding sites. X-axis shows the genomic feature. Y-axis shows the percentage of SVs. Top 30 TFs that bind to these SV overlapping sites the most are listed on a separate horizontal bar chart, listing TFs on the y-axis and its frequency on the x-axis. **d**, Bar chart showing the composition of SV types in each public database. Databases are on the x-axis and count is on the y-axis. The four standard SV types (INS, DEL, DUP, and INV) are color-coded, and all other types are binned together into the “Other” group colored in gray.

**Extended Data Fig. 4 | ASE and eQTL analysis for HLA genes. a**, Bubble plot showing the results of ASE and eQTL in midbrain (left) and MTG (right). Significant SV-gene pairs in HLA locus (chr6: 28,510,120-33,480,577) are plotted. Information about SV-gene pairs is summarized in row names in a 5-column format (chromosome, position, SV type, SV length, and gene name). Cell types tested are specified on the x-axis. Adjusted p-value is shown by the bubble size (-log_10_ scale) and effect size is shown by the color. Significant associations have black outline. Same SV affecting multiple genes is shown by a dotted horizontal line. SV-gene pairs on the y-axis are sorted by chromosome and position. **b,c**, Dot plots showing the expression of HLA genes in midbrain, **b**, and MTG, **c**. Dot size shows percent of cells expressing the gene, and color shows scaled average expression.

**Extended Data Fig. 5 | ASE and eQTL analysis of SV-gene pairs identified in only one of the analyses. a-b**, Bubble plots showing the results of ASE and eQTL. Brain region and type of test performed are indicated at the top. **a** includes only the SV-gene pairs significant in midbrain and **b** includes only the SV-gene pairs significant in MTG. Information about SV-gene pairs is summarized in row names in a 5-column format (chromosome, position, SV type, SV length, and gene name). Cell types tested are specified on the x-axis. Adjusted p-value is shown by the bubble size (-log_10_ scale) and effect size is shown by the color. Significant associations have black outline. Same SV affecting multiple genes is shown by the dotted horizontal line. SV-gene pairs on the y-axis are sorted by chromosome and position.

**Extended Data Fig. 6 | Examples of ASE and eQTL gene expression. a**-**g,i,k**, Box plots showing the pseudobulk expression of SV-eQTL significant target genes from snRNA-seq data across cell types in MTG, **a**-**c,f,g**, and midbrain, **d,e**,**i,k**. SV-gene pair and cell types are indicated at the right and top, respectively. Individuals are grouped by SV genotype (x-axis) and their averaged expression values (y-axis) are plotted. Individual data points are color-coded by the diagnosis. **a**, Genotypes listed as ./. indicate the SV caller did not report a genotype, though they are likely 0/0. The slash indicates that the genotype is unphased, because it is an inversion, which did not have Kanpig genotyping applied. **h,j**, Box plots showing the proportion of phased UMIs mapping to the alternate allele (x-axis) across cell types (y-axis). SV-gene pair is indicated at the right. MTG, **h**, and midbrain, **j**, examples are shown. Asterisks show statistical significance as in Fig. 5.

**Extended Data Fig. 7 | Examples of ASE and eQTL with confidence intervals. a**-**l**, Dot and whisker plots showing effect size (x-axis) with confidence intervals across cell types (y-axis). ASE shown on the left and eQTL on the right. Both brain regions, midbrain (top) and MTG (bottom), are shown. SV-gene pair is indicated at the top. Color represents significance, where red indicates adjusted p-value < 0.05 and black is not significant.

**Extended Data Fig. 8 | ASE analysis with hybrid selected snRNA-seq data. a-b**, Bubble plots showing the results of ASE analysis on targeted snRNA-seq data in midbrain, **a**, and MTG, **b**. Only significant SV-gene pairs are plotted. Information about SV-gene pairs is summarized in row names in a 5-column format (chromosome, position, SV type, SV length, and gene name). Cell types tested are specified on the x-axis. Adjusted p-value is shown by the bubble size (-log_10_ scale) and effect size is shown by the color. Significant associations have black outline. Same SV affecting multiple genes is shown by the dotted horizontal line. SV-gene pairs on the y-axis are sorted by chromosome and position.

**Extended Data Fig. 9 | Additional ASE and eQTL analysis. a**,**b**, Scatter plots comparing the effect sizes from ASE with and without hybrid selection analyses across different cell types in midbrain, **a**, and MTG, **b**. Effect sizes from ASE with the selection data are shown on the x-axis, and those from ASE with non-selection data are shown on the y-axis. Dots are color-coded by significance in the two methods. Only SV-gene pairs significant in at least one method are plotted. **c**, Bar charts showing the number of unique significant SVs from ASE analysis in the midbrain (black) and MTG (red). The number of significant SVs is on the x-axis and cell types are listed on the y-axis. ASE without hybrid selection (left) and ASE with hybrid selection (right) shown. **d**, Heatmap showing the correlation between ASE or eQTL effect size and SV properties across cell types for each SV type in MTG and midbrain. Brain region and the analysis type are indicated on the left. SV types are specified at the top. SV properties include SV length and minor allele frequency (y-axis). Cell types are shown on the x-axis. Statistical significance was indicated by asterisks based on adjusted p-value thresholds: * Padj <= 0.05, ** Padj <= 0.01, *** Padj <= 0.001, and **** Padj <= 0.0001. No asterisk means Padj > 0.05. **e**,**f** Bar charts showing the potential reason for the cell type-specific eQTL and ASE signals in the MTG, **e**, and midbrain, **f**. Number of significant SV-gene pairs are plotted on the x-axis, and cell types are on the y-axis. Plots are divided into three columns based on what is driving the effect: low expression (bottom), low expression and cell number (middle), and potential regulatory specificity (top). Number of SV-gene pairs from ASE analysis (red) and eQTL analysis (grey) are shown.

**Extended Data Fig. 10 | Genomic features near SV examples. a,** Two SVs with significant ASE and eQTL effects on *DDX55*. Center track shows gene models and SVs. The upper inset panel focuses on the 974 bp insertion. Single base position (purple rectangle) where this insertion occurs is shown at the top of this panel. The track below SVs shows chromatin a state segmentation (wgEncodeBroadHmm) from ENCODE that was converted via liftOver to hg38. Chromatin state groups are indicated on the left (also color-coded by the state). The tracks below show accessible regions from PsychENCODE across different cell types (brainSCOPE). Cell types are indicated on the left and in the description within the tracks. The bottom track shows repeat structures from RepeatMasker and Dfam databases. The top insert shows the structure of the inserted sequence. The bottom inset panel focuses on the 108 bp deletion. The deletion is shown at the top (purple rectangle). The same set of tracks are shown for this deletion. **b**, Intronic region of *BIN3* locus with ASE/eQTL significant 330 bp insertion. Tracks are as explained in **a**.

## Supplementary Figures

**Supplementary Fig. 1 | SV caller performance. a**, Bar chart showing the genotype agreement between two SV caller combinations across SV types. Different caller combinations are color-coded. SV types are shown on the x-axis. Percent of genotype agreement is shown on the y-axis. Note only one caller combination is shown for the insertions because Cue2 does not call insertions by design. **b**, Bar charts showing Kanpig benchmarking for synthetic datasets. Synthetic datasets include three different sequence repeat contexts: no repeat (left), short repeat (middle), and segmental duplication (right). Simulated SVs are further categorized by SV type (INS, DEL, DUP, and INV) and size (bp, x-axis). Genotype accuracy based on the ground truth is shown on the y-axis. SV types are color-coded. **c**, Bar charts showing VaPoR benchmarking for synthetic inversions, as in **b**.

**Supplementary Fig. 2 | Cell type composition and eGene expression in snRNA-seq data. a**,**b**, Heatmaps showing the snRNA-seq expression of eGenes across cell types in MTG, **a**, and midbrain, **b**. Rows show eGenes, shown with hierarchical clustering using Euclidean distance, and columns show cell types. Expression is scaled for each gene and z-score is shown. **c**,**d**, Bar charts showing the composition of cell types in MTG, **c**, and midbrain, **d**. Cell types are color-coded and specified on the x-axis. Y-axis shows number of cells.

## Supplementary Tables

Supplementary Table 1. PD5D cohort

Supplementary Table 2. DNA QC

Supplementary Table 3. HiFi read QC

Supplementary Table 4. SV QC

Supplementary Table 5. SV annotation

Supplementary Table 6. Public SV catalogs

Supplementary Table 7. SVs for functional analysis

Supplementary Table 8. SV-eQTL results

Supplementary Table 9. SV-ASE results

Supplementary Table 10. SV-ASE hybrid selection summary

Supplementary Table 11. Key resource table

Supplementary Table 12. Comparison of eQTLs with and without population stratification

## Supplementary Note 1

Using simulated HiFi reads and SVs has a few advantages: (1) we know the ground truth to assess the genotyper accurately; (2) we can simulate SVs in various sequence contexts including non-repeat, short repeats, and segmental duplications (repeat size >1 kb). (3) SVs of different sizes can be simulated. Genotype concordance (number of correctly predicted SV genotypes over the total) of Kanpig prediction to the ground truth for synthetic datasets is presented in Supplementary Fig. 1b. Kanpig’s accuracy varied with sequence context, SV type, and SV size. First, the genotyping accuracy in non-repeat and short repeat regions were very similar (maximum concordance 99.7%) but dropped sharply in segmental duplications (maximum concordance 51.3%). We anticipated this difficulty in segmental duplications and SVs overlapping these regions were removed during quality control. Second, SV type also has notable effect. In both non-repeat and short repeat synthetic datasets, while the median concordance for insertions, deletions, and duplications was > 97%, that of inversions was < 1%. As such, we tested VaPoR^38^, which is better suited for more complex rearrangements (Supplementary Fig. 1c). However, benchmarking on synthetic dataset suggested that the genotype concordance between VaPoR prediction and the ground truth in the repeat context is at best 65% in short repeat and 50% in segmental duplications (Supplementary Fig. 1c). Thus, we used caller predicted genotype calls for inversions. Third, SV size affected Kanpig genotype accuracy, as expected^37^, with lower accuracy for SVs > 10kb for insertions and deletions and for duplications > 1kb in both non-repeat and short repeats.

Therefore, for subsequent analysis, we decided to only include SVs that we have confidence in assigning accurate genotypes. We define Kanpig confident ranges in non-repeat and short-repeat: insertions of size within 50 bp – 10 kb (minimum genotype concordance = 93.3%), deletions of size within 50 bp – 10 kb (minimum genotype concordance = 89.6%), and duplications of size within 50 bp – 1 kb (minimum genotype concordance = 97.7%). Kanpig failed to accurately assign genotypes for inversions of all sizes in any sequence repeat contexts (maximum genotype concordance = 2.7%). By contrast, VaPoR was able to assign inversion genotypes accurately in non-repeat regions (minimum genotype concordance = 90%). However, this concordance drops in repeat regions (maximum genotype concordance = 65%). As such, we decided to use the caller predicted genotypes for inversions and include all sizes.

## Supplementary Note 2

For the eQTL analyses presented in this study, we omitted covariates from population stratification (genotype PCs) as described in the Methods to be consistent with the prior SNP-based eQTL results generated by our team. To assess the effects of population stratification though, we compared eQTL results with and without six genotype PCs as additional covariates (Supplementary Table 12). The results were comparable: 755 SV-gene pairs were significant in both analyses, 211 were significant only with population stratification, and 219 were significant only without population stratification.

## Notes

### Competing Interest Statement

The authors have declared no competing interest.

### Summary of Updates

Supplementary Table 1 has been revised -- changing ages 90 and above to "90+" and removing dates from DX summary field.

