## Extended Data Figures for "Integrating Long-Read Structural Variant Analysis with single-nucleus RNA-seq to Elucidate Gene Expression Effects in Disease"

### Extended Data Fig. 1

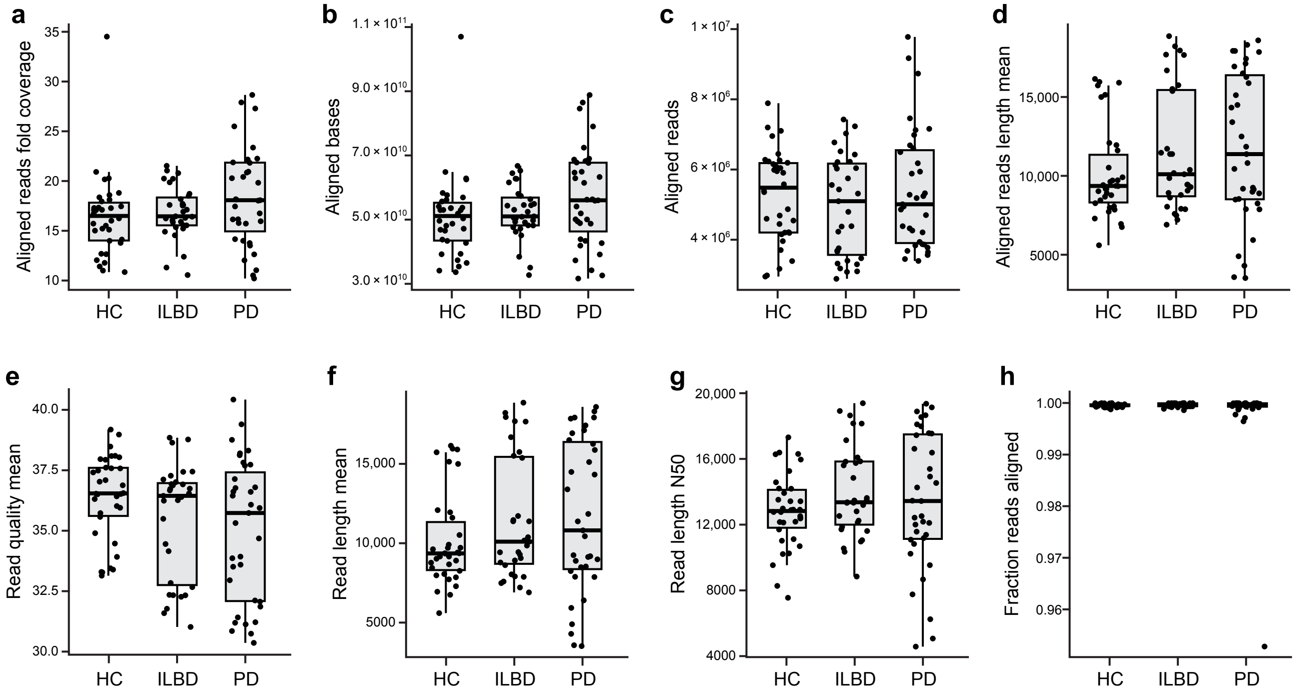

### Extended Data Fig. 2

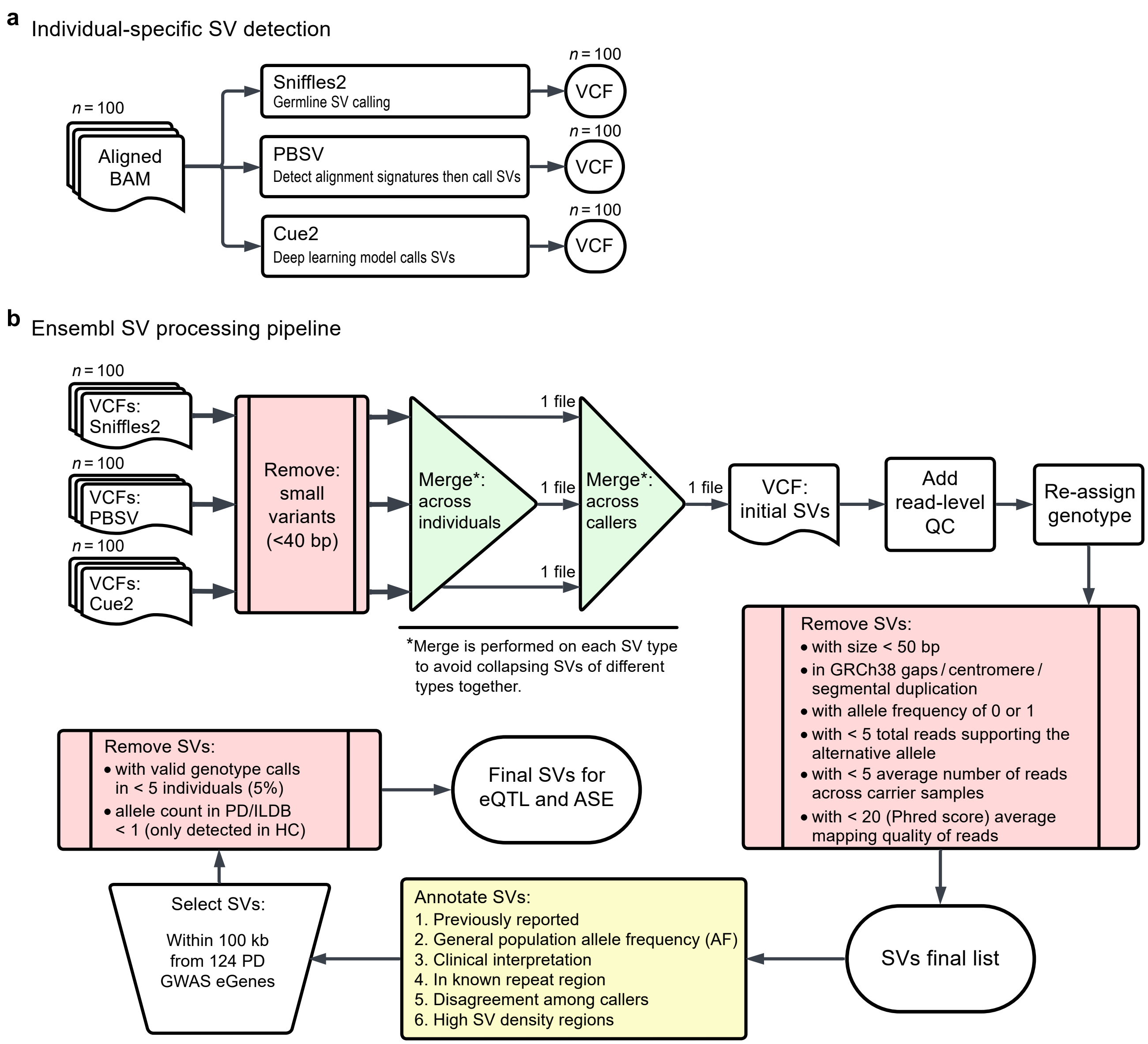

### Extended Data Fig. 3

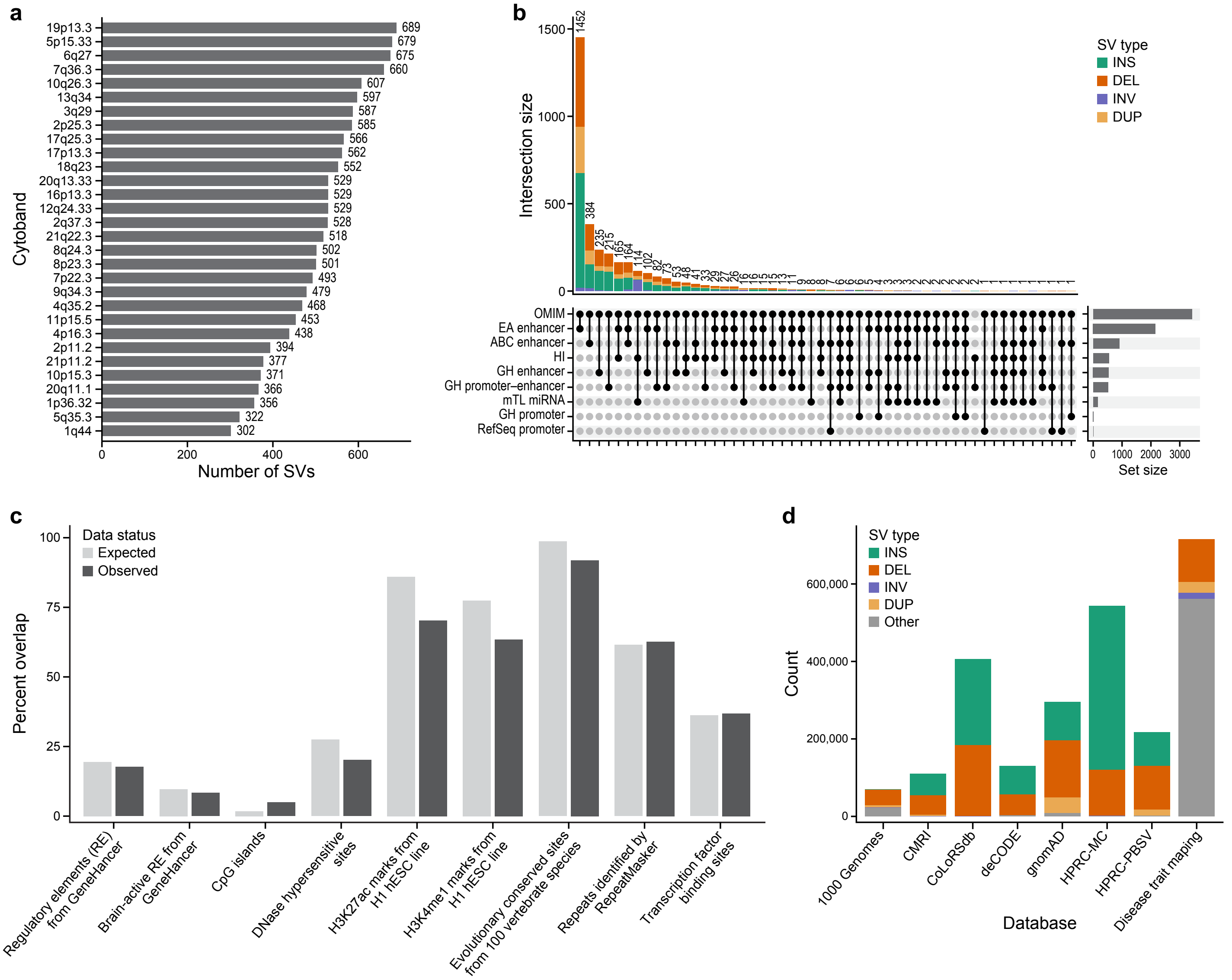

### Extended Data Fig. 4

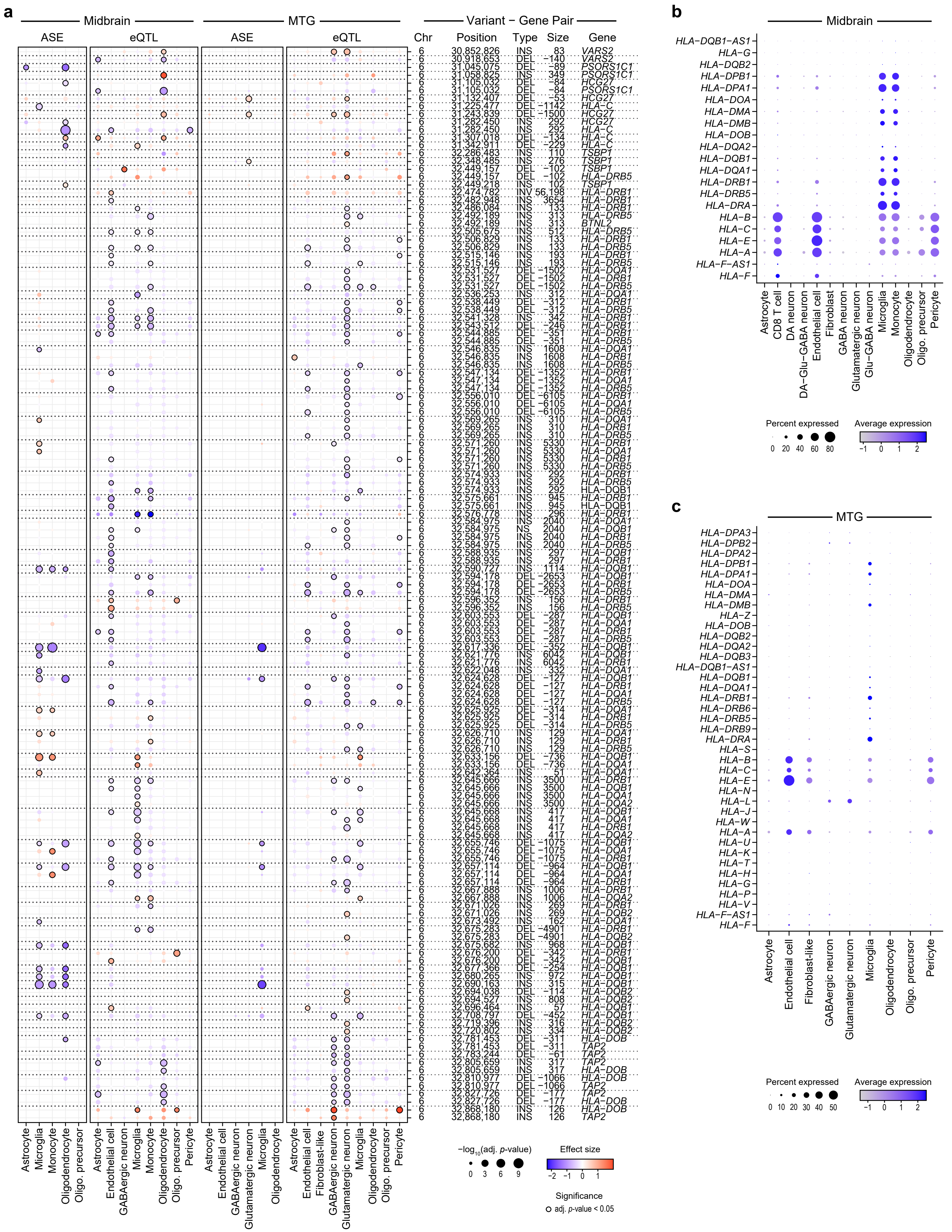

### Extended Data Fig. 5

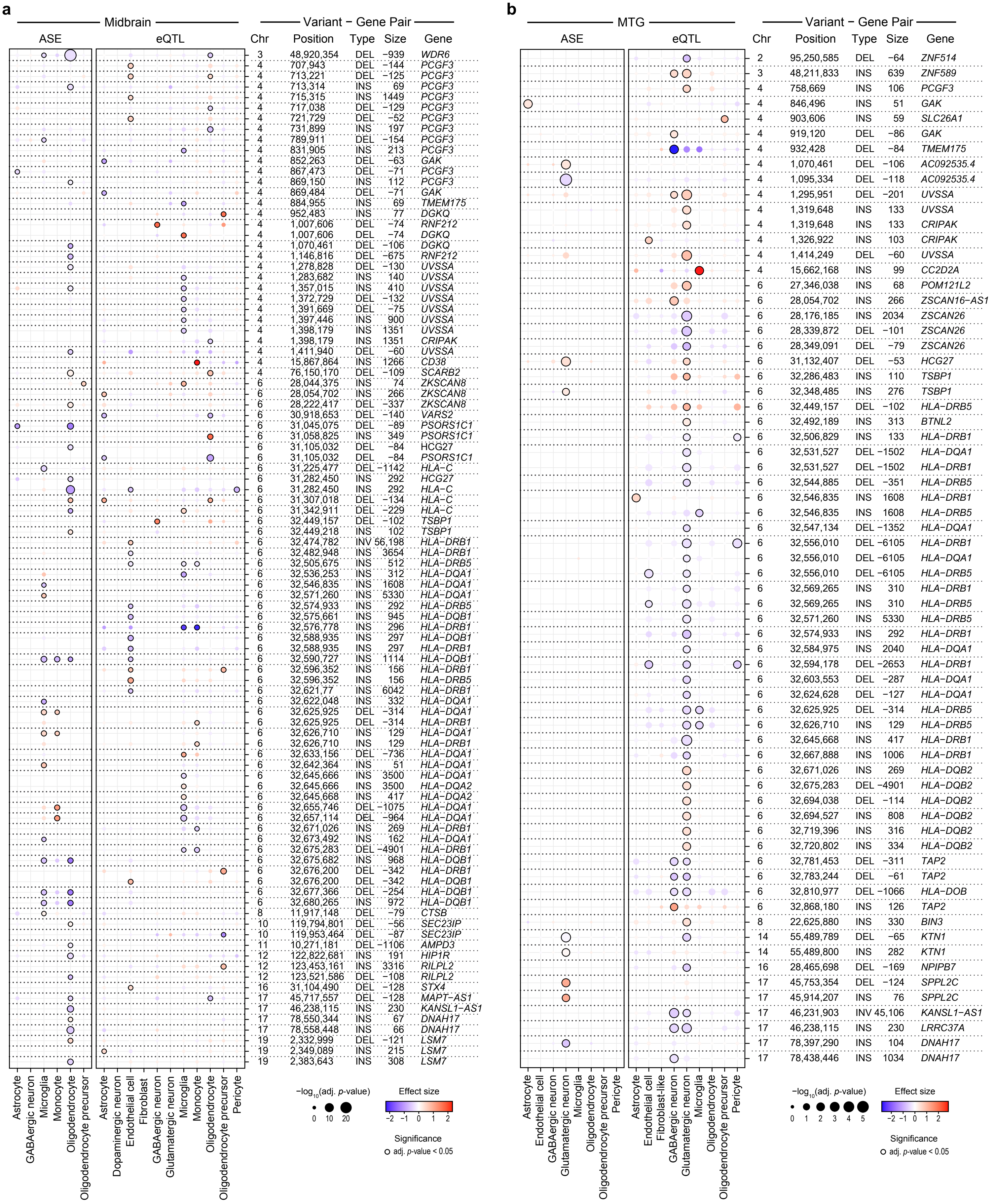

### Extended Data Fig. 6

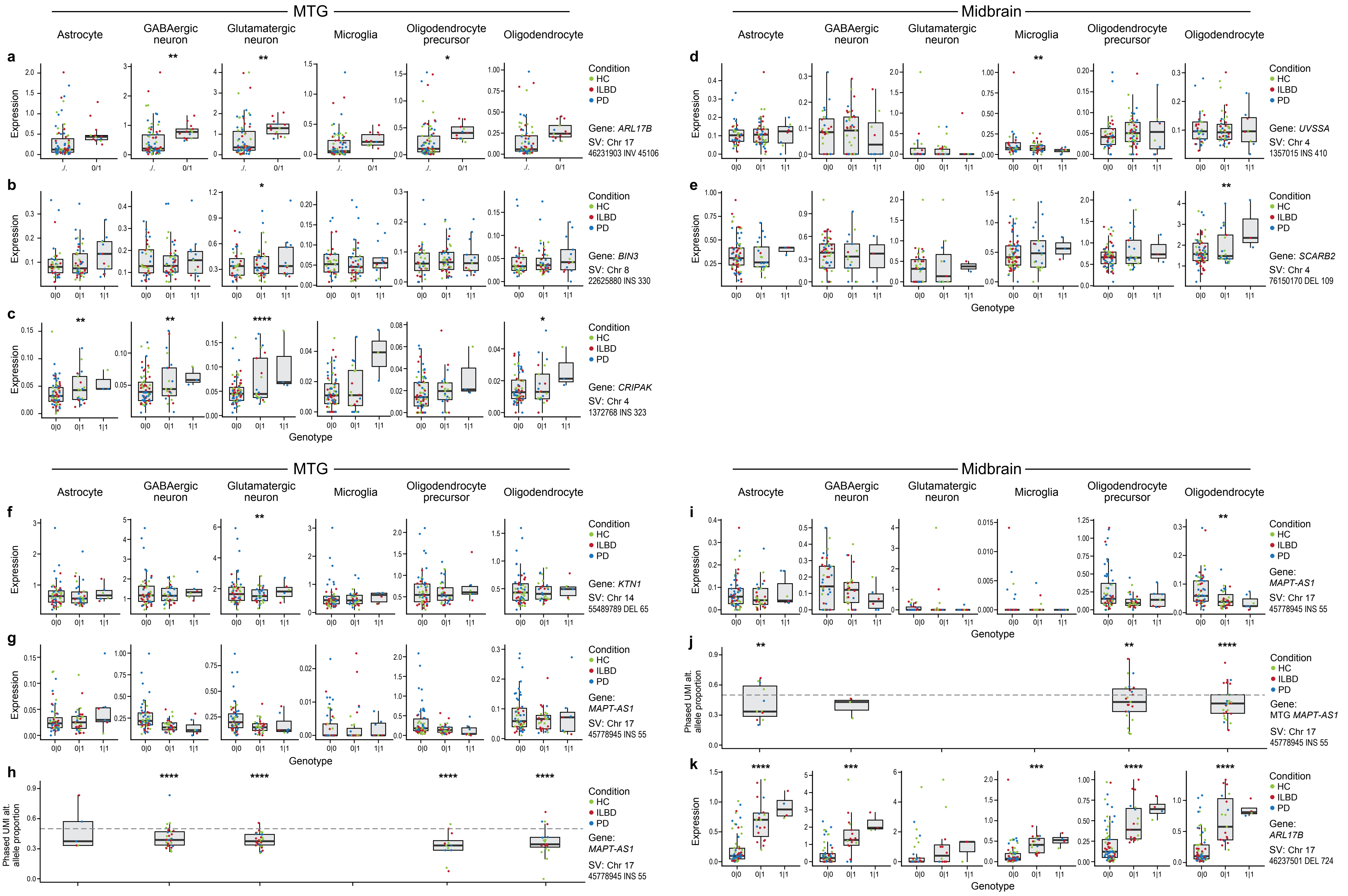

### Extended Data Fig. 7

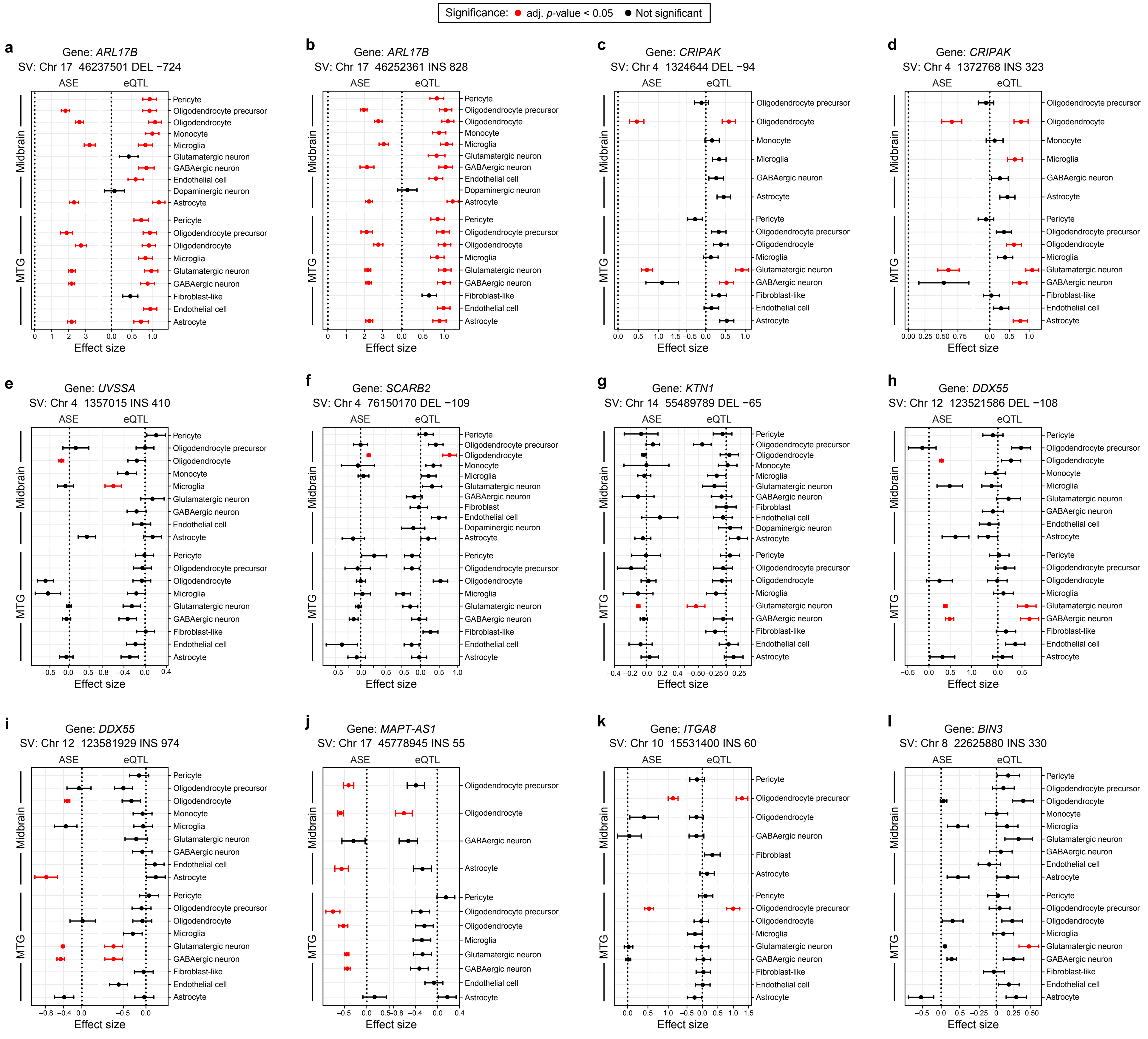

### Extended Data Fig. 8

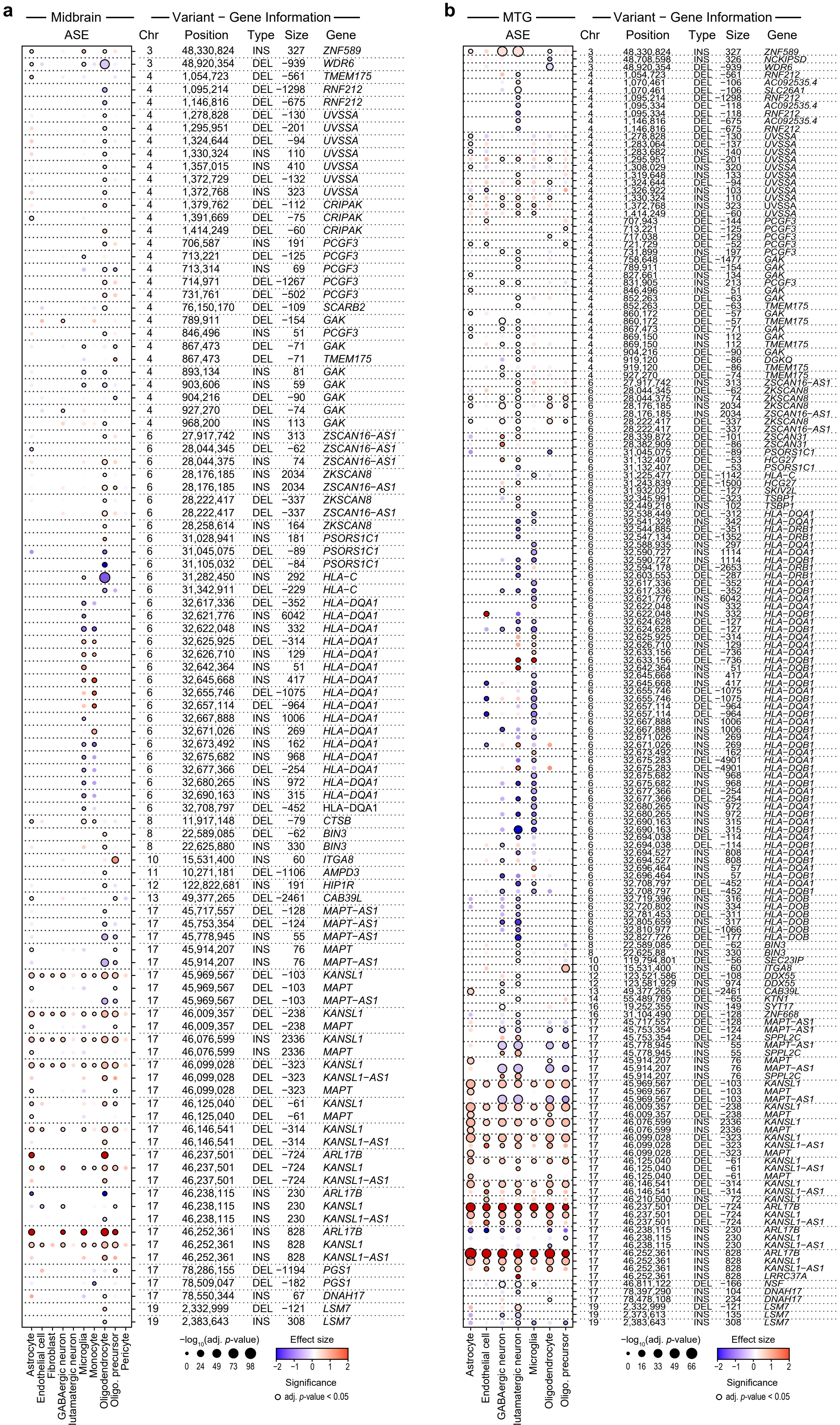

### Extended Data Fig. 9

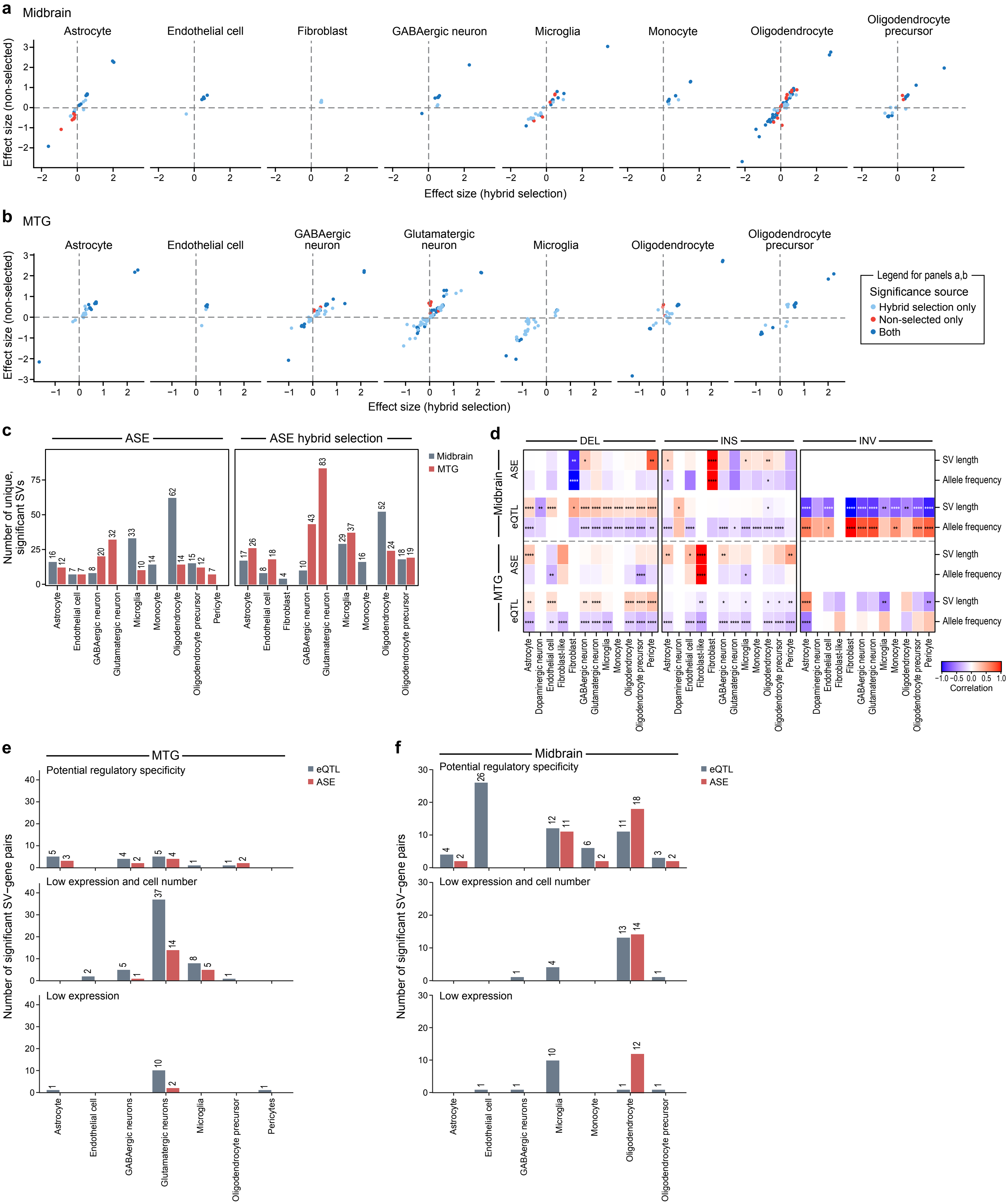

### Extended Data Fig. 10

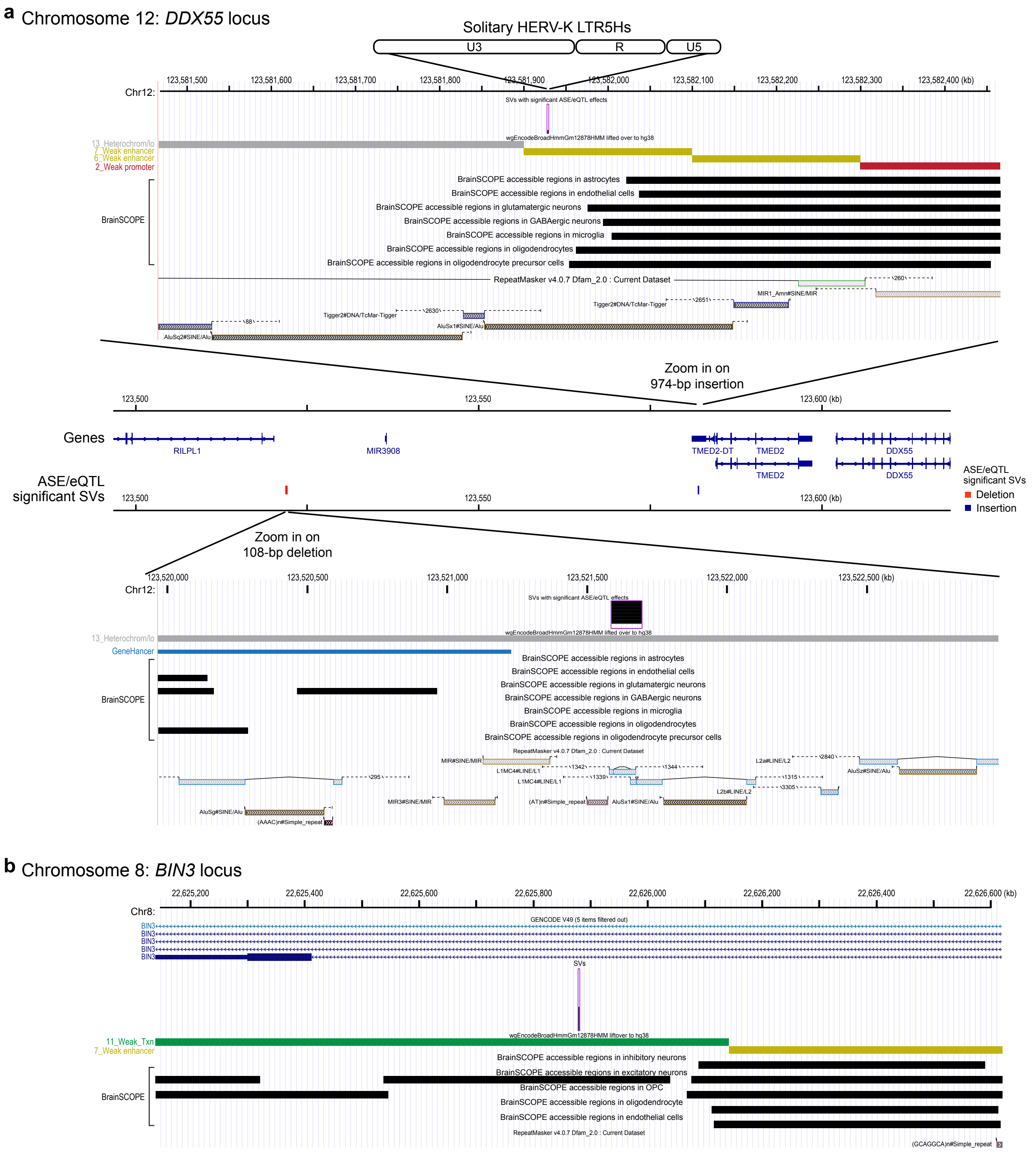
