## Supplementary figures and images for "Integrating Long-Read Structural Variant Analysis with single-nucleus RNA-seq to Elucidate Gene Expression Effects in Disease"

### Supplementary Data Figures

#### Supplemental Fig. 1

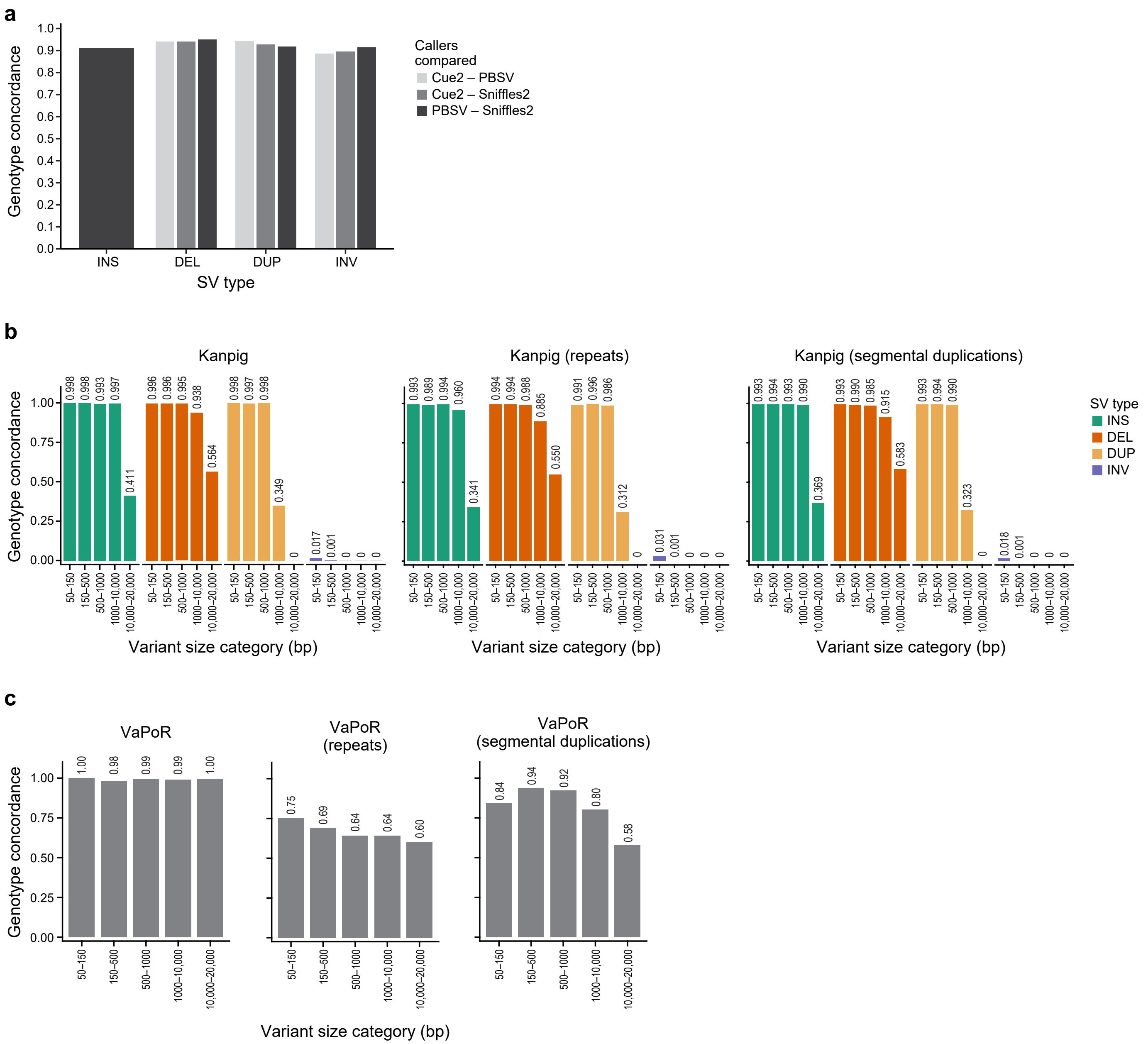

Supplemental Fig. 2

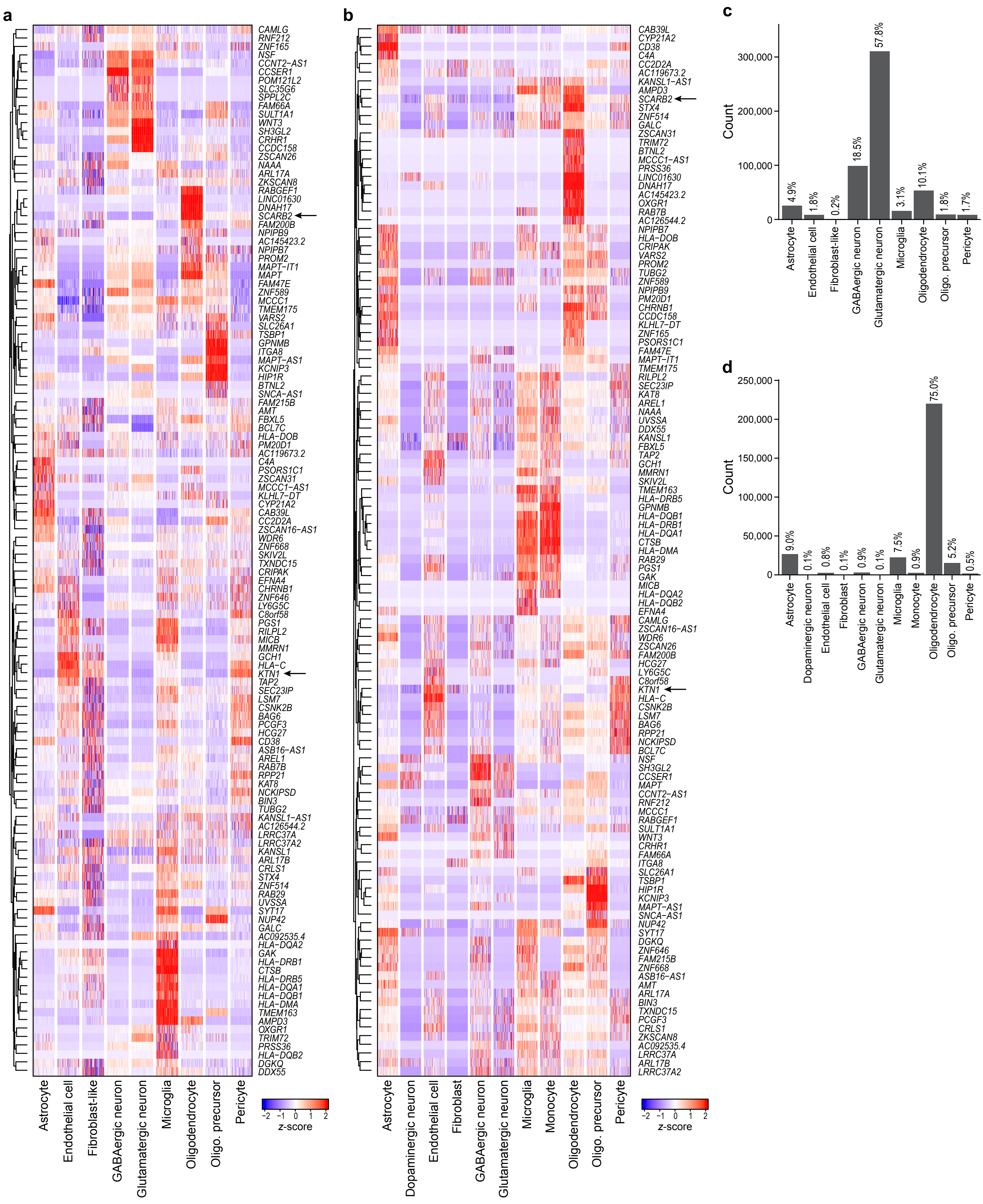
